# Emergence of a Cell-Guided Multivalent Ligand of Enzymes on Cancer Cells Triggered by Click Reaction between Hetero Nano-Assemblies

**DOI:** 10.1101/2025.04.24.649770

**Authors:** Rentaro Sakamoto, Yuki Koba, Masahiko Nakamoto, Tatsuya Fukuta, Kazunori Kadota, Michiya Matsusaki

## Abstract

Stimuli-responsive nanomaterials with multivalent ligands have attracted significant attention in the context of cancer chemotherapy and imaging. However, challenges remain such as non-selective stimuli response and/or existence of triggers in healthy regions in addition to the intrinsic heterogeneity of cancer that causes insufficient target recognition. Inspired by the expression of precise and diverse functions of biological machineries triggered by specific protein-protein complexation and conformational change, we report an artificial system where a bioorthogonal click reaction between hetero-nano-assemblies triggers their complexation and conformational changes, resulting in emergence of multivalent ligands for cancer-associated enzymes: carbonic anhydrase IX (CAIX). We also demonstrated that the multivalent ligands selectively inhibited the proliferation of cancer cells overexpressing CAIX under hypoxic conditions. Additionally, the click reaction between nano-assemblies in the presence of target cells provided a higher efficacy of the emerged multivalent ligands than that pre-formed in the absence of cells. Our study provides a basis for the development of multivalent ligands displaying adaptive binding interfaces for target cancer cells with high selectivity and affinity to thus potentially overcome tumor heterogeneity.

## Introduction

Nanomaterials such as polymers, nanoparticles, and self-assemblies have been extensively studied in the fields of cancer therapy and/or imaging due to their multifunctionalities and/or enhanced permeation and retention effects.^[1-3]^ Engineering stimuli-responsive nanomateri-als that change their structure in response to pH, enzymes, and/or light and release anticancer drugs and/or promote their cancer cell affinity represent a promising strategy for the improvement of cancer selectivity.^[4-13]^ Kim et al. reported an approach in which L-histidine-based polymeric micelles that release anticancer drugs triggered by endosomal acidic environments.^[9]^ Molla et al. demon-strated target protein-triggered disassembly of amphiphilic polypeptide-based nanoaggregates that displayed ligands on the hydrophilic face and the consequent release of payloads.^[10]^ A stimuli-triggered decrease or increase in the size of materials has also been exploited to control their penetration or retention within the dense tumor stroma.^[11-13]^

Employing multivalent interactions by functionalizing nanomaterials with multiple ligands is also effective for the selective recognition of target proteins and cells.^[14-23]^ In particular, multivalent ligands that exhibit switching affinity for target proteins/cells in response to stimuli have gained much attention.^[24-29]^ Dreher et al. reported the temperature-triggered self-assembly of polypeptides into micelles with multivalent peptide ligands at their termini, which actualized temperature-responsive cellular uptake.^[24]^ Kohane et al. reported light-triggered activation of multivalent recognition of cells by polymer self-assemblies decorated with photocaged cell-penetrating peptides for the intravenous treatment of choroidal neo-vascularization.^[27]^ However, despite significant efforts, nanomaterials-based cancer targeting remains a challenge due to undesired activation by nonspecific responses and/or the existence of triggers in healthy regions. Moreover, recent studies have revealed that the intrinsic heterogeneity of cancer causes an insufficient response at the tumor site.^[30-32]^ The spatiotemporal heterogeneity of target enzymes/proteins on cancer cells would cause the difficult prediction of optimal chemical/physical structures of multivalent ligands prior to administration as well, because the density and/or spatial relation of multivalent ligands and receptors strongly affect their avidity.^[33, 34]^ These drawbacks highlight the importance of specifically triggered and target cell-guided *in situ* ligand multimerization for the selective and adaptive targeting of cancer cells.

Click chemistry, established by Sharpless and Meldal et al., is used in a variety of chemical reactions due to its high orthogonality, high yield, and fast reaction rate.^[35-37]^ Click reactions have further evolved into ligand synthesis in the presence of target enzymes/proteins. “target-guided synthesis” of small molecule-based enzyme inhibitors ^[38-41]^ and peptide-based protein capture reagent ^[42]^ have demonstrated efficient lead discovery. Since Bertozzi et al. introduced Cu-free strain-promoted azide-alkyne cycloaddition (SPAAC), bioorthogonal click chemistry has become a valuable tool for understanding and regulating biological processes.^[43-45]^ Bioorthogonal click chemistry has allowed the development of specific and chemically triggered payload release systems for cancer therapy and imaging.^[46-50]^ Robillard et al. developed a bioorthogonal elimination reaction that enabled the instantaneous, self-immolative, and traceless release of a substance from trans-cyclooctene following tetrazine ligation.^[46]^ Based on this concept of “click to release”, they demonstrated the release of anticancer drugs from a non-internalizing antibody-drug conjugate triggered by a subsequently administrated chemical activator in murine tumor models.^[47]^ Porte et al. reported tumor imaging of xenograft mice using fluorophore-loaded micelles that disassembled via a sequential activation process consist-ing of an enzymatic reaction and strain-promoted iminosydnone-cycloalkyne cycloaddition.^[48]^

Bioorthogonal click chemistry has also facilitated the engineering of chemically triggered *in situ* functionalization of nanomaterial-based systems. A reversible click reaction between boronic acid and salicyl hydroxamate has been employed as a strategy for post-polymerization modification to prepare polymer-drug conjugates that self-assemble into nanostructures, accomplishing the release of conjugated drugs in response to versatile stimu-li in the tumor microenvironment such as acidic, oxidation, and/or reduction conditions.^[49]^ Wang et al. constructed *in situ* self-assembled aggregates on the cancer cell membrane using SPAAC between peptides that function as target recognition and hydrophobic motifs, respectively, thus enabling membrane disruption and enhancing the chemodrug sensitivity of renal cell carcinoma.^[51]^ Tanaka et al. employed SPAAC for *in situ* ligation of two ligands based on saccharides and peptides with different affinities for corresponding receptors, and this significantly improved cell recognition ability.^[52]^ However, an artificial system in which the *in situ* multimerization of ligands is triggered by a biorthogonal click reaction between nanomaterials to intervene in the cellular function of interest has not yet been reported.

In signal transductions in living systems, diverse, adaptive and precise functions emerge by *in situ* specific protein-protein complexation and subsequent conformational changes even though the number of available proteins and their elemental functions are limited. ^[53-57]^ For instance, interactions between intrinsically disordered regions of B-cell lymphoma-2 (Bcl-2) family proteins induce their conformational changes, which regulates Bcl-2 function and the subsequent apoptotic pathway.^[58]^ Taking inspiration from these molecular machineries, we are the first to report an artificial system in which *in situ* SPAAC between hetero-nano-assemblies triggers their complexation, conformational changes, and the emergence of multivalent ligands for a cancer-associated enzyme to inhibit cancer cell proliferation (Figure 1). The results also revealed that *in situ* emergence of multivalent interactions guided by the target cells improved their proliferative inhibitory efficacy. Our study would be a strategy for creating multivalent ligands that display an adaptive binding interface for target cancer cells with high selectivity and affinity that could overcome tumor heterogeneity. Our concept would form the basis for *in situ* functionalization of nanomedicines as well.

**Figure 1.**
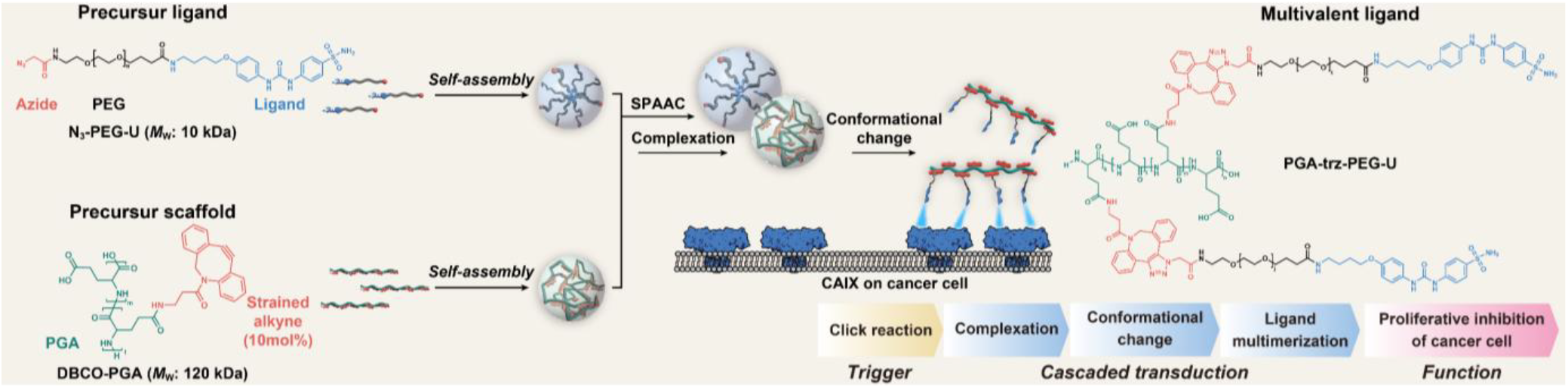
Conceptual illustration of this study.

## Results and Discussion

### Synthesis and characterization of N_3_-PEG-U as precursor ligand

A hypoxia-induced transmembrane enzyme, carbonic anhydrase IX (CAIX), was selected as the target enzyme, as it plays a vital role in cancer cell metabolism and is a promising target for cancer therapy.^[28, 29, 51, 59-61]^ As a precursor of ligand, a polyethylene glycol (PEG)-based nano-assembly with CAIX ligand (U-104) and azide (N_3_) group was synthesized in three steps (Supporting Information 2 and 3). First, U-104 was introduced into alpha-t-butyloxycarbonylamino-omega-carboxy succinimidyl ester PEG (Boc-PEG-NHS; average molecular weight, 10 kDa), resulting in Boc-PEG-U. Subsequently, the Boc protection was removed under acidic conditions (NH_2_-PEG-U), and N_3_-AcOH was introduced at the amino terminus, resulting in N_3_-PEG-U with either a monovalent U-104 or N_3_ group at each terminus (Figure 2A and S-1 and Scheme S-2). The obtained N_3_-PEG-U was well-dispersed in D-PBS due to its amphiphilic nature, although U-104s exhibited low solubility when the concentration of the U-104 group was fixed (Figure S-2). Dynamic light scattering (DLS) measurement revealed that N_3_-PEG-U formed self-assembly in the aqueous solution and exhibited a hydrodynamic diameter of 159 ± 3 nm (Figure 2B). We also confirmed that the introduction of U-104 at the termini of Boc-PEG-NHS resulted in self-assembly and that subsequent Boc deprotection and azidation at the remaining termini did not strongly affect its size. These results indicate that intermolecular interactions such as hydrogen bonding, hydrophobic interactions, and/or interactions between ureido-substituted benzene sulfonamide of U-104s ^[62]^ are the driving forces of self-assembly. Thus, N_3_ groups are exposed at the surface of the assemblies. Circular dichroism spectrometry also indicated the right-hand helicity of U-104s in the self-assemblies (Figure S-3). Pyrene fluorescence titrations ^[63]^ and DLS measurements ^[64]^ were used to determine the critical self-assembly concentration (CSC) of N_3_-PEG-U at 10 mM and 3.2 mM (Figure 2C and S-4).

**Figure 2.**
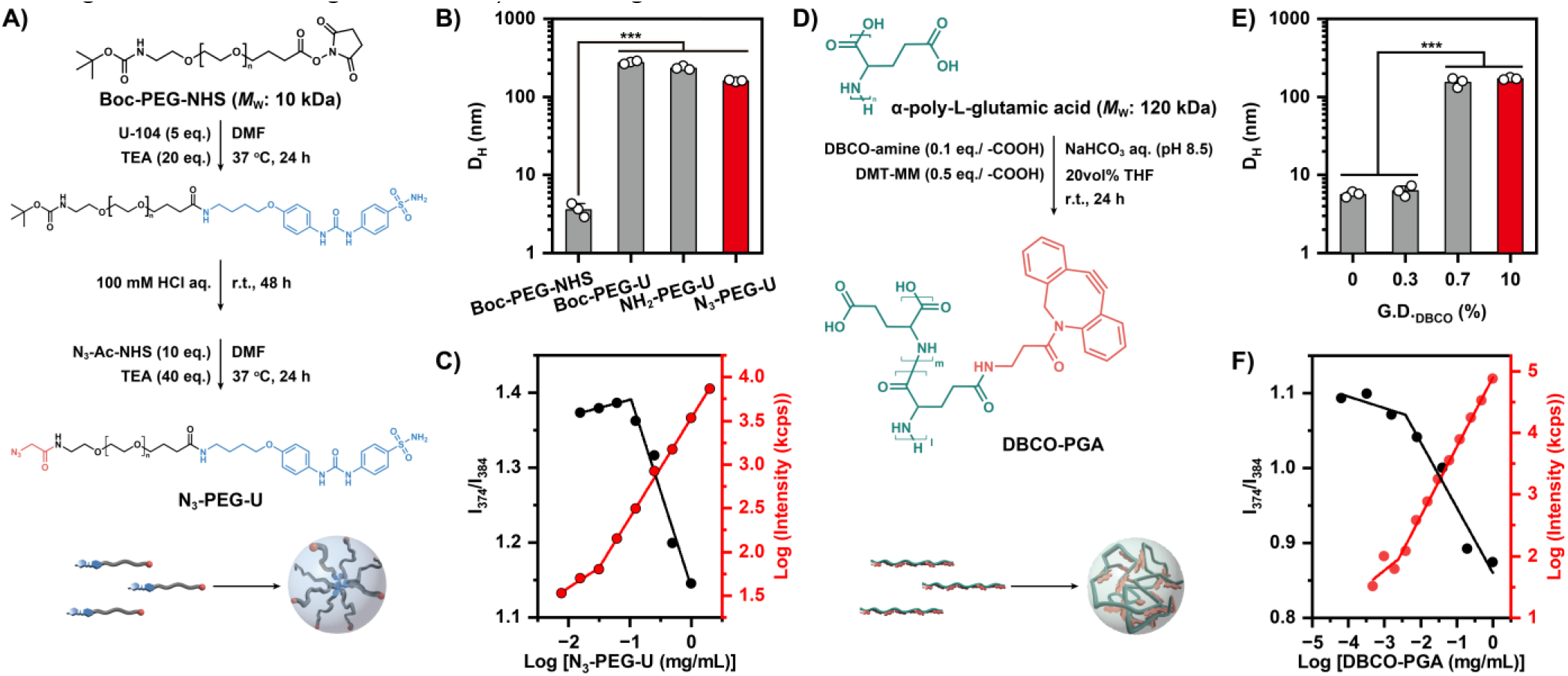
A) Synthesis scheme, B) volume average hydrodynamic diameter and C) CSC determination of precursor ligand: N3-PEG-U. D) Synthesis scheme, E) volume average hydrodynamic diameter and F) CSC determination of precursor scaffold: DBCO-PGA. Statis-tical significance was determined using one-way ANOVA followed by Tukey’s post-test (n = 3) (*n*.*s*.: not significant, * *p* < 0.05, ** *p* < 0.01, *** *p* < 0.001). Data are mean ± SD.

### Synthesis and characterization of DBCO-PGA as precursor scaffold

As a precursor of the scaffold of a multivalent ligand, a-poly-L-glutamic acid (PGA)-based self-assembly with multivalent strained alkyne was synthesized by introducing dibenzocyclooctyne-amine (DBCO-amine) to sidechains of PGA with an average molecular weight of 120 kDa (Figure 2D and Supporting Information 5).The improved dispersity of amphiphilic DBCO-PGA compared to that of DBCO-amine was confirmed in D-PBS at a fixed concentration of the DBCO group (Figure S-6). DLS revealed a drastic increase in size when the introduction amount of DBCO was 0.7 mol% or more, indicating that polymers formed self-assemblies by inter-polymer multipoint hydrophobic and/or π-π interaction (Figure 2E). DBCO-PGA with 10 mol% of grafting degree and a hydro-dynamic diameter of 170 ± 5 nm was used in further experiments. The CSC values of DBCO-PGA obtained from pyrene fluorescence titrations and DLS measurements were 0.032 and 0.020 mM, respectively (Figure 2F and S-7)

### SPAAC triggered the conformational transformation of nano-assemblies

Next, we investigated the impact of SPAAC between two nanoassemblies, N_3_-PEG-U and DBCO-PGA, on the size and morphology of click product, PGA-trz-PEG-U. To that end, the hydrodynamic diameter of a mixture of N_3_-PEG-U ([N_3_] = 150 μM) and DBCO-PGA ([DBCO] = 200 μM) in D-PBS at 37 °C was observed as a function of time (Figure 3A). The hydrodynamic diameters of either N_3_-PEG-U alone or DBCO-PGA alone were approximately 100 nm and 180 nm, and no size change was observed for both nanoassemblies after 24 h (Figure S-8). When the two nanoassemblies were mixed, an obvious size reduction was observed within 30 min (Figure 3A), resulting in a peak with a 30 nm diameter after 6 h accompanied with the reaction progress (Figure 3B and Figure S9). We also confirmed that PGA-trz-PEG-U did not show obvious CSC values (Figure S-10). These results indicate that the SPAAC reaction between N_3_-PEG-U and DBCO-PGA induced complexation and conformational changes, ultimately yielding PGA-trz-PEG-U with a reduced size. SPAAC-triggered conformational changes were confirmed using transmission electron microscopy (TEM) observations. N_3_-PEG-U and DBCO-PGA exhibited a spherical shape, with circularities of 0.73 and 0.77, respectively. However, PGA-trz-PEG-U exhibited a fragmented shape with a significantly lower circularity of 0.68 and a broad distribution (Figure 3C and 3D). Small angle X-ray scattering (SAXS) profile of the mixture of N_3_-PEG-U and DBCO-PGA also indicated time dependent increase of scattering derived from click products with smaller diameter than the precursors; N_3_-PEG-U alone and DBCO-PGA alone (Figure S-11). Taken together, we concluded that the conformational change in the nano-assemblies is triggered by SPAAC. Reportedly, the self-assembling property of polymers is affected not only by the chemical structure of assembling groups but also by various factors, such as sequence, molecular weight and morphology of precursor polymers.^[65, 66]^ Thus it is more likely that SPAAC reaction reduces the stability of each assembly by providing electrostatic/steric repulsion to N_3_-PEG-U and steric repulsion/hydrophilization to DBCO-PGA (Figure 3E). Calculation of the LogP values ^[67]^ of analogs of the

**Figure 3.**
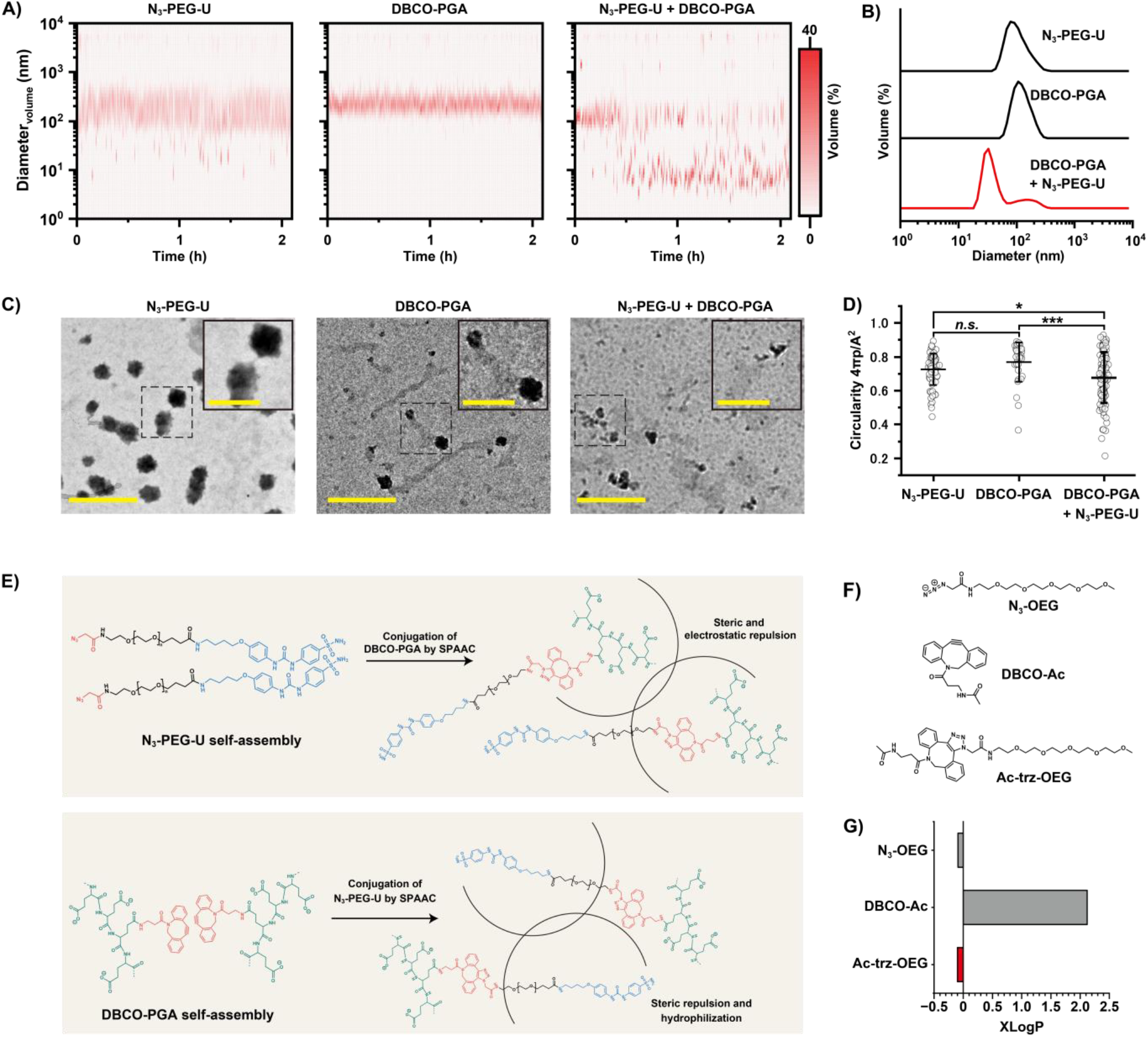
A) Size distribution of N_3_-PEG-U, DBCO-PGA, and PGA-trz-PEG-U in D-PBS at pH 7.4 at 37 °C measured by DLS. B) Time courses of N_3_-PEG-U, DBCO-PGA, and PGA-trz-PEG-U. C) TEM images (the insets show a magnified images, Scale bar: 200 nm and 50 nm (inset)) and D) circularity of N_3_-PEG-U, DBCO-PGA and mixture. E) Proposed perturbation to N_3_-PEG-U and DBCO-PGA upon SPAAC. F) Chemical structure and G) calculated LogP values of DBCO-Ac, N_3_-OEG and Ac-trz-OEG. Statistical significance was determined using one-way ANOVA followed by Tukey’s post-test (n > 30) (*n*.*s*.: not significant, * *p* < 0.05, ** *p* < 0.01, *** *p* < 0.001). Data are mean ± SD.

DBCO side chain (DBCO-acetate), oligoethylene glycol azide (N_3_-OEG), and their SPAAC product (Ac-trz-OEG) also sup-ports the local hydrophilization effect on DBCO groups upon SPAAC (Figure 3F and 3G). For further understanding, the SPAAC reaction kinetics were analyzed (Figure 4). To elucidate the influence of self-assembly on the reaction, the small-molecule substrates N_3_-AcOH and DBCO-amine were used for comparison in addition to N_3_-PEG-U and DBCO-PGA. Reactions were monitored based on a decrease of UV absorption derived from alkyne groups (λ= 309 nm) at a fixed concentration of N_3_ groups and DBCO groups at 50 mM in D-PBS at 37 °C (Figure 4A, 4B and S-12-15). The reaction rate constants were obtained assuming second-order reaction kinetics (Figure 4C, Figure S-16-19 and Table S1). The reaction rate constant between N_3_-PEG-U and DBCO-amine was comparable to that between the small-molecule substrates N_3_-AcOH and DBCO-amine. Hence, it is likely that the N_3_ group was exposed on the surface of the N_3_-PEG-U nanoassembly. In contrast, the reaction rate constant between DBCO-PGA and N_3_-AcOH was considerably smaller than that between small-molecule substrates, indicating buried DBCO groups in the DBCO-PGA nanoassemblies. These results are consistent with our discussion of the formation mechanism of the aforementioned self-assemblies. The reaction rate constant between N_3_-PEG-U and DBCO-PGA was moderate; however, it was higher than that between DBCO-PGA and N_3_-AcOH. This result suggests that the conjugation of N_3_-PEG-U with highly hydrophilic and flexible PEG chains to DBCO-PGA (i.e., PEGlyation) disaggregated the DBCO-PGA nano-assembly and allowed DBCOs to be exposed to bulk aqueous solution during the reaction (Figure 4D). This is consistent with the reaction-triggered conformational changes of the assemblies observed by DLS and TEM.

**Figure 4.**
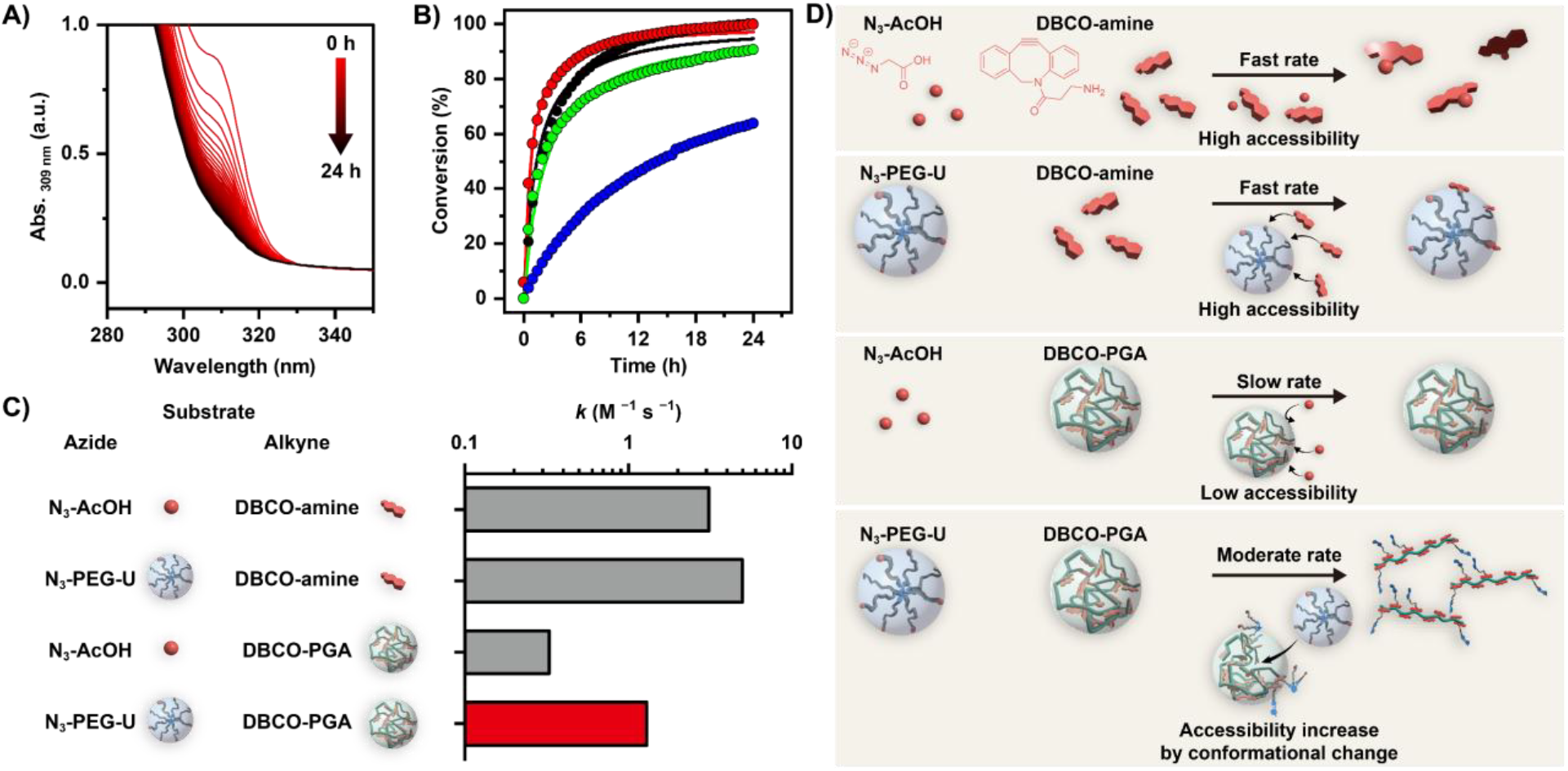
A) Typical UV-vis spectra for click reaction between N_3_-PEG-U and DBCO-PGA. B) Time courses of conversion for SPAAC. Filled dots and solid lines show the data and fitting curve, respectively (black: N_3_-AcOH and DBCO-amine, red: N_3_-PEG-U and DBCO-amine, blue: N_3_-AcOH and DBCO-PGA and green: N--PEG-U and DBCO-PGA). The fitting parameter was shown at Table S-1. C) Reaction rate constants of tested azide and DBCO substrates. D) Schematic illustration of SPAAC between tested substrates.

### The emergence of multivalent interactions with target enzymes triggered by SPAAC

To validate the click reaction-triggered multivalent recognition of CAIX, the binding of N_3_-PEG-U, DBCO-PGA, and PGA-trz-PEG-U to the CAIX surface was analyzed using a 27 MHz quartz crystal microbalance (QCM) (Figure 5A). Streptavidin was immobilized on a biotin-modified gold substrate via biotin-avidin interactions (Figure S-20). Subsequently, biotin-tagged CAIX (CAIX-Fc-biotin) was immobilized on a streptavidin monolayer, resulting in an oriented CAIX surface (Figure S-21). Casein was used as a blocking agent to prevent nonspecific interactions. PGA-trz-PEG-U with a multivalency of 70 was prepared by incubating the mixture of N_3_-PEG-U and DBCO-PGA in D-PBS at 37°C for 20 hours before the binding analysis (Supporting information 9). The frequency decrease with the addition of PGA-trz-PEG-U was significantly larger than that with the addition of N_3_-PEG-U and DBCO-PGA at the same concentration of N_3_ and DBCO (500 nM). The -ΔF values of N_3_-PEG-U, DBCO-PGA and PGA-trz-PEG-U were 42 ± 24 Hz, 44 ± 19 Hz, and 130 ± 10 Hz, respectively (Figure 5B and S-22-24). We further assessed the contribution of the multivalency of the ligands to their binding affinities and amounts on the CAIX surface (Figure 5C). To this end, we synthesized PGA-trz-PEG-U with multivalencies of 15, 30, and 70 (Figure S-25-28). The saturated binding amount and apparent dissociation equilibrium constants (*K*_d_) of PGA-trz-PEG-U toward the CAIX surface were quantified assuming Langmuir-type binding (Figure 5D). When multivalency was 30 or less increasing the multivalency tended to increase both the binding amount and affinity of PGA-trz-PEG-U for the CAIX-immobilized surface. *K*_d_ of PGA-trz-PEG-Us was significantly lower than that of the small molecule U-104 ^[68]^, indicating enhanced affinity by multivalent interactions. The binding amount and affinity were saturated or slightly depleted when the multivalency was 70 (N_3_ / DBCO was 1 / 1), and this may indicate that excess multivalency resulted in increased steric hindrance or ligand self-association. These results conclude that multivalent binding of PGA-trz-PEG-U to CAIXs occurs via SPAAC-triggered cascade multimerization and conformational changes in the assemblies.

**Figure 5.**
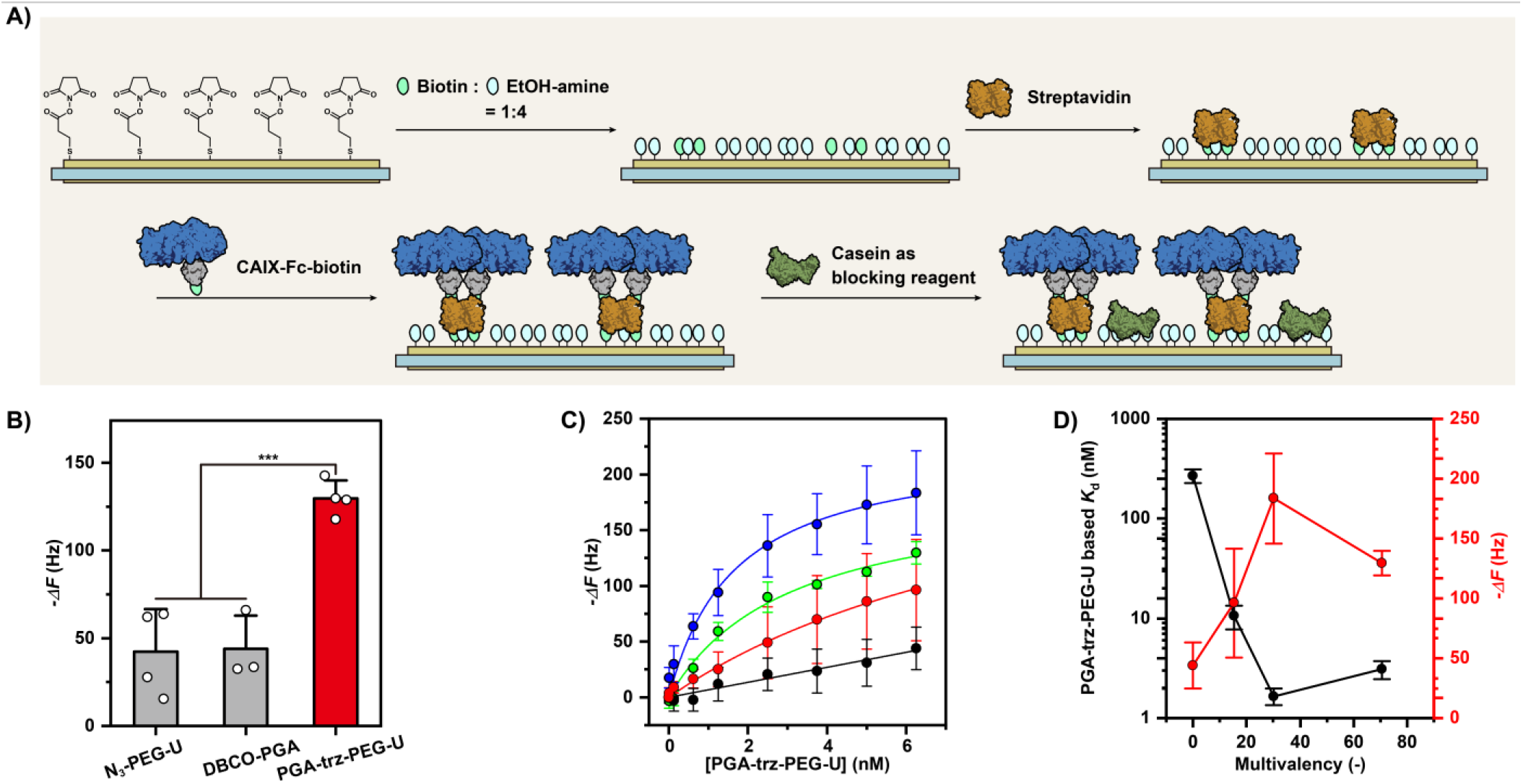
A) Experimental set up of QCM measurement. B) Binding of N_3_-PEG-U, DBCO-PGA and PGA-trz-PEG-U at fixed concentration ([U] = [DBCO] = 500 nM). C) Binding isotherm of PGA-trz-PEG-U with various multivalency on CAIX immobilized surface (black: multivalency of 0, red: multivalency of 15, blue: multivalency of 30, and green: multivalency of 70). Note that PGA-trz-PEG-U with multivalency of 0 is intact DBCO-PGA. (D) Binding amount and affinity of PGA-trz-PEG-U with various multivalency on CAIX immobilized surface. Statistical significance was determined using one-way ANOVA followed by Tukey’s post-test (n ≧ 3) (*n*.*s*.: not significant, * *p* < 0.05, ** *p* < 0.01, *** *p* < 0.001). Data are mean ± SD.

### The emergence of multivalent interactions with target enzymes triggered by SPAAC

The proliferative inhibitory effects of N_3_-PEG-U, DBCO-PGA, and PGA-trz-PEG-U on the human breast cancer cell line MDA-MB-231 under hypoxic (1% O_2_) and slightly acidic conditions (pH 6.8) in which CAIX is overexpressed were evaluated using a water-soluble tetrazolium salt assay (WST-8 assay) (Figure 6A). We first confirmed that the precursors, N_3_-PEG-U and DBCO-PGA, did not affect cell proliferation at the tested concentrations. Additionally, the small-molecule ligand (U-104) exerted an obvious inhibitory effect on cell proliferation at the same ligand concentration as that of N_3_-PEG-U, indicating the inactivation of CAIX recognition by self-assembly. Next, the emergence of the SPAAC-triggered inhibition of cancer cell proliferation between precursors was demonstrated by the co-addition of N_3_-PEG-U and DBCO-PGA to cells. Although neither N_3_-PEG-U alone nor DBCO-PGA alone caused any inhibition, the combination of N_3_-PEG-U and DBCO-PGA significantly inhibited cell proliferation (Figure 6A and S-29). This result concludes that the emerged multivalent ligands of CAIX inhibited the proliferation of cancer cells overexpressing CAIX. It was also confirmed that the addition of the polymer alone or in combination did not inhibit cell proliferation under normoxic or neutral conditions (pH 7.4) where CAIX was not overexpressed.^[29, 59]^ This result indicates the selective potency of PGA-trz-PEG-U for cancer cells that overexpress CAIX under hypoxic conditions. The effect of the multivalency of PGA-trz-PEG-U on the inhibition of proliferation was further evaluated at a fixed PGA-trz-PEG-U concentration. The efficacy of the proliferative inhibition increased with increasing multivalency (Figure 6B). Subsequently, the effect of PGA-trz-PEG-U concentration on proliferative inhibition was evaluated at a fixed multivalency of 40. The efficacy increased with increasing polymer concentrations (Figure 6C). Finally, we assessed the impact of *in situ* ligand synthesis on its efficacy. To that end, The efficacy of *in situ* PGA-trz-PEG-U synthesized in the presence of cells and *ex situ* PGA-trz-PEG-U pre-synthesized in the absence of cells was compared (Figures 6D and E). Both ligands were synthesized using a DBCO-to-azide molar ratio of 2:1; the resulting ligand multivalency was 40, and 50% of the DBCO groups remained. We confirmed that the SPAAC reaction was complete within 1 hour in cell culture medium under the provided conditions during *in situ* PGA-trz-PEG-U synthesis (Figure S30) and the multivalency was the same as that of the *ex situ* ligand (Figure S31). Despite having the same multivalency, *in situ* PGA-trz-PEG-U exerted significantly higher proliferative inhibition than *ex situ* PGA-trz-PEG-U (Figure 6D), indicating the higher target cell affinity of *in situ* ligand than *ex situ* one. These results imply that the positioning of ligands on the scaffold was guided by target enzymes on cells during multimerization, accompanied by conformational changes (Figure 6E).

**Figure 6.**
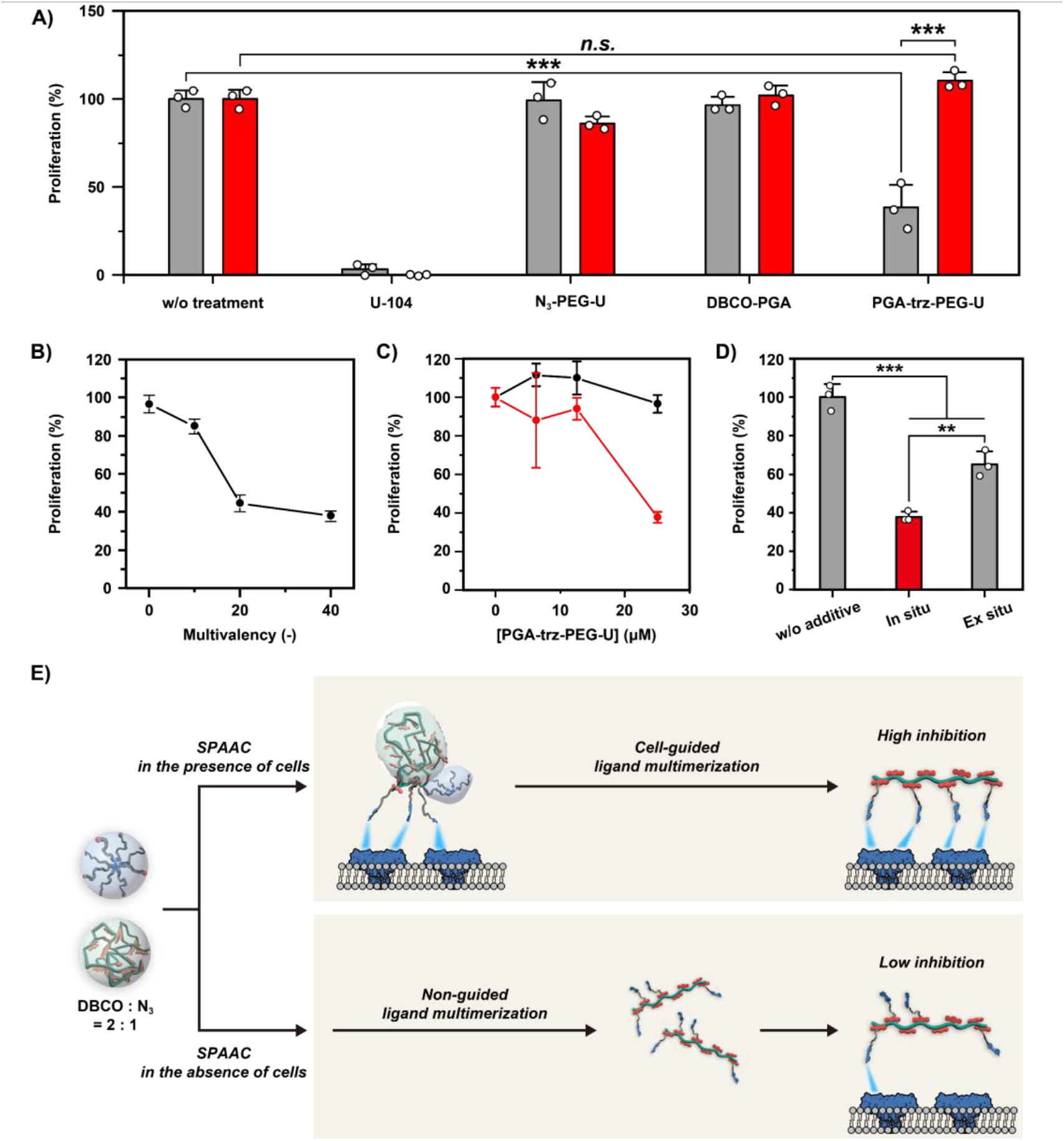
A) Cell proliferation of MDA-MB-231 treated with U-104 ([U] = 1000 μM), N_3_-PEG-U ([N_3_-PEG-U] and [N_3_] = 1000 μM) and/or DBCO-PGA ([DBCO-PGA] = 25 μM and [DBCO] = 2000 μM) (black: hypoxia and pH 6.8 and red: normoxia and pH 7.4). B) Effect of the multivalency of PGA-trz-PEG-U ([PGA-trz-PEG-U] = 25 μM). C) Effect of the concentration of PGA-trz-PEG-U (black: multivalency is 0, red: multivalency is 40). Note that PGA-trz-PEG-U with multivalency of 0 is intact DBCO-PGA. D) Effect of *in situ* emergence of PGA-trz-PEG-U on cell proliferative inhibition. E) Schematic illustration of non-guided (left) and cell-guided ligand multimerization (right). Statistical significance was determined using one-way ANOVA followed by Tukey’s post-test (n = 3) (*n*.*s*.: not significant, * *p* < 0.05, ** *p* < 0.01, *** *p* < 0.001). Data are mean ± SD.

We also visualized the polymers that bound to MDA-MB-231 cells using fluorescence-labeled N_3_-PEG-U (N_3_-PEG-U-Cy5) and DBCO-PGA (DBCO-PGA-Cy3) (Figure S-32 and S-33). The Cy5 fluorescence intensities of cancer cells treated with both N_3_-PEG-U and DBCO-PGA were significantly higher than those treated with N_3_-PEG-U alone when the polymer concentration was low (the concentrations of the N_3_ and DBCO groups were 40 and 80 μM, respectively) (Figure 7A, B, S-34, and S-35). This result suggests the activation of the recognition of cancer cells via SPAAC. However, the Cy5 fluorescent intensities of cancer cells treated with N_3_-PEG-U at an N_3_ concentration of 200 µM were not significantly different from that of co-treated cancer cells, and this is likely due to non-specific internalization at higher polymer concentrations. The Cy3 fluorescence intensities of cancer cells treated with DBCO-PGA alone were comparable to or slightly higher than those of co-treated cells, and this was independent of concentration (Figure 7C, S-36, and S-37), indicating non-specific adsorption of DBCO-PGA due to its hydrophobic nature. It should be noted that non-specific internalization/adsorption of N_3_-PEG-U/DBCO-PGA would not affect cell proliferation. The Cy3 fluorescence intensity of the co-treated cells could also be lowered via energy transfer to Cy5 that was closely introduced by SPAAC. To evaluate the Förster resonance energy transfer (FRET) of the co-treated cells, FRET factors were compared (Figure 7D, S-38, and S-39). The FRET factor of the co-treated cells was significantly higher than that of cells treated with DBCO-PGA alone, suggesting the adsorption of PGA-trz-PEG-U on the cells.

**Figure 7.**
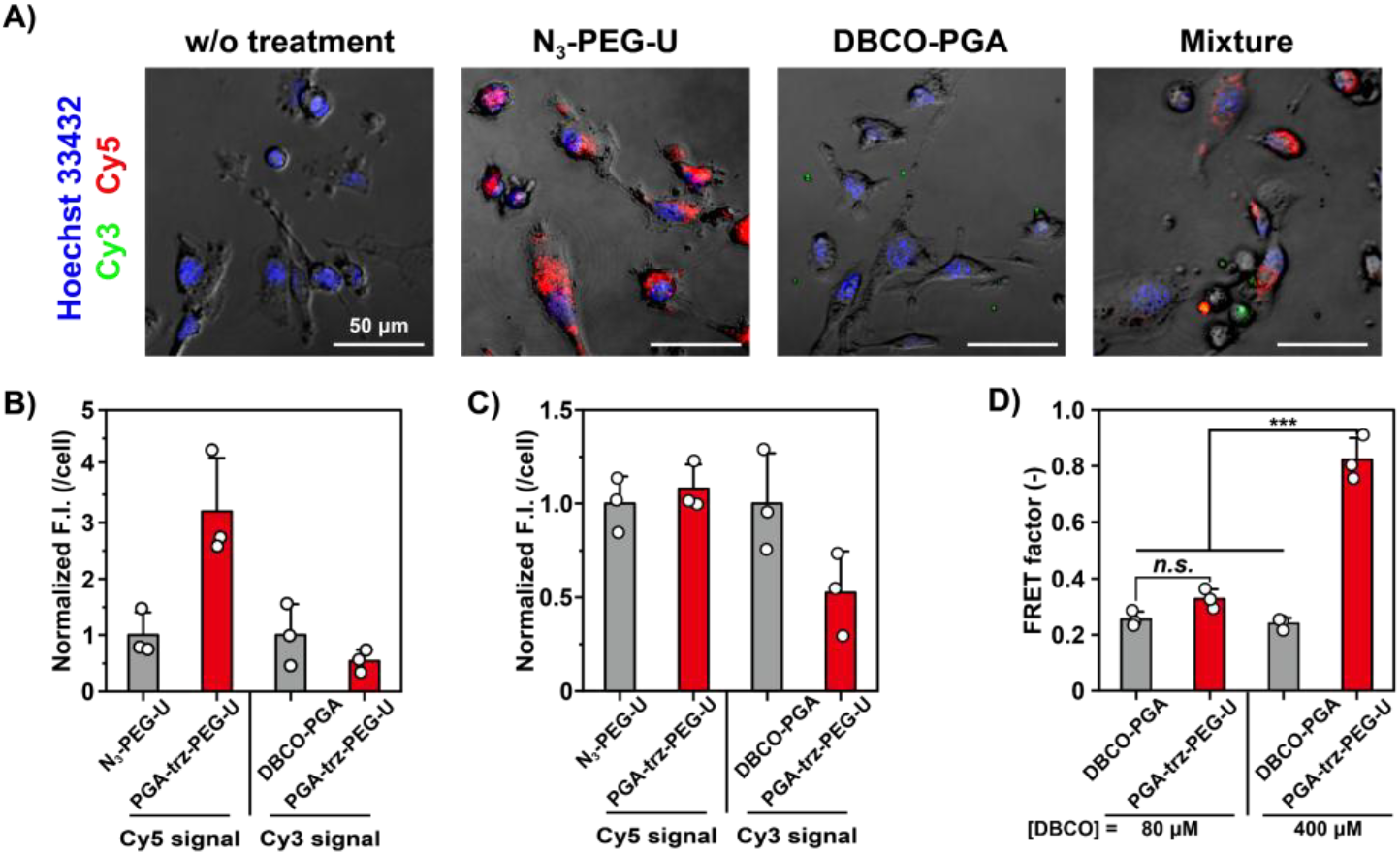
A) CLSM images of MDA-MB-231 treated with N_3_-PEG-U ([N_3_-PEG-U] and [N_3_]= 40 μM) and/or DBCO-PGA ([DBCO-PGA] = 1 μM and [DBCO] = 80 μM). The scale bar is 50 mm. B) The fluorescent intensities of Cy5 and C) Cy3. D) FRET factor of MDA-MB-231 treated fluorescence labelled polymers. Statistical significance was determined using one-way ANOVA followed by Tukey’s post-test (n = 3) (*n*.*s*.: not significant, * *p* < 0.05, ** *p* < 0.01, *** *p* < 0.001). Data are mean ± SD.

## Conclusion

Inspired by the molecular machinery involved in signal transduction triggered by specific protein-protein complexation, we first reported an artificial system in which SPAAC between hetero-nano-assemblies triggers the emergence of multivalent ligands for cancer-associated enzymes. SPAAC between the nano-assemblies as precursor scaffolds and ligands resulted in their complexation and conformational changes that emerged as multivalent ligands with high affinity for CAIX-immobilized surfaces. The proliferative inhibition of cancer cells overexpressing CAIX using this system was also demonstrated. Additionally, SPAAC between precursors in the presence of target cells provided a higher efficacy of multivalent ligands than that preformed in the absence of cells, indicating the cell-guided emergence of multivalent ligands. Our study provides a basis for the development of multivalent ligands displaying adaptive binding interfaces for target cancer cells with high selectivity and affinity that could overcome tumor heterogeneity.

## Supporting Information

The authors have cited additional references within the Supporting Information. ^[69, 70]^

## Acknowledgements

We would like to thank Prof. K. Mitsuoka of Research Center for Ultra-High Voltage Electron Microscopy, Osaka University, for assistance in the TEM observations. We are also grateful to Anton Paar, for their support with SAXS experiments and the analysis.

## Entry for the Table of Contents

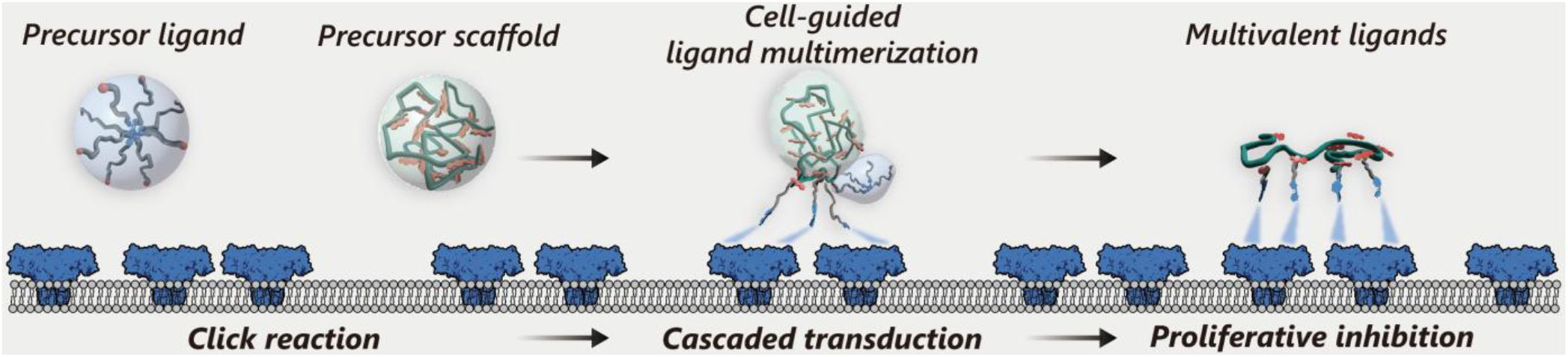

### Cell-guided emergence of multivalent ligands for cancer cells

Click reaction between nano-assemblies led to complexation and conformational changes, resulting in the emergence of multivalent ligands with high affinity for cancer cells. The multivalent ligands synthesized in the presence of cancer cells more effectively inhibited their proliferation than those synthesized in their absence, indicating cell-guided effect.

