## Supporting information for "Emergence of a Cell-Guided Multivalent Ligand of Enzymes on Cancer Cells Triggered by Click Reaction between Hetero Nano-Assemblies"

Table of contents

1. Material

2. Synthesis of CAIX inhibitor U-104

3. Synthesis of $N_3$-PEG-U

4. Evaluation of physico-chemical properties for $N_3$-PEG-U

5. Synthesis of DBCO-PGA

6. Evaluation of physico-chemical properties for DBCO-PGA

7. Change of physico-chemical properties of nano-assemblies via click reactio

8. Reaction kinetics evaluation of click reaction

9. Binding assay by quartz crystal microbalance

10. Cell experiments

### 1. Material

#### Reagents

Unless otherwise noted, all other commercially available reagents and solvents were used without further purification. Tetrahydrofuran (207-17765), 5 mol/L sodium hydroxide solution (1310-73-2), Potassium hydrogen sulfate (7646-93-7), Toluene super dehydrated (108-88-3), Acetonitrile super dehydrated (75-05-8), 4 M Hydrogen chloride・ethyl acetate solution (083-10405), *N,N*-dimethylformamide (045-32365), 5 mol/L hydrochloric acid (7647-01-0), Sodium carbonate (199-01585), Sodium hydrogen carbonate (191-01305), Sulfuric Acid (7664-93-9), *N*-hydroxy succinimide (6066-82-6), 2-amino ethanol (141-43-5), Streptavidin (9013-20-1) and Casein, from milk (9000-71-9) were obtained from Wako, Japan. Methyl 4-hydroxybenzoate (99-76-3), Triphenylphosphine (603-35-0), *N, N*- diisopropylethylamine (7087-68-5), Diphenyl phosphoryl azide (26386-88-9), Sulfanilamide (63-74-1), Triethylamine (121-44-8), Pyrene (129-00-0), Azide acetic acid (18523-48-3), 1-ethyl-3-(3-dimethylaminopropyl) carbodiimide (25952-53-8), Biotin-PEG2-amine (121-44-8) and 4-dimethylaminopyridine (1122-58-3) were obtained from Tokyo Chemical Industry Co., Ltd., Japan. Dulbecco' s Phosphate Buffered Saline (07269-84), Ethanol (64-17-5), Dulbecco's modified Eagle's medium (08458-45), Antibiotic-Antimycotic Mixed Stock Solution (100x) (02892-54), Cell Count Reagent SF (07553-44) and Hoechst 33342 Solution (1 mg/mL) (23491-52-3) were obtained from nacalai tesque, Japan. Diisopropyl azodicarboxylate (2446-

83-5), Chloroform-d (865-49-6), Dimethyl sulfoxide-D6 (2206-617-0), Deuterium oxide (7789-20-0), Dibenzocyclooctyne-amine (1255942-06-3) and 3,3'-dithiodipropionic acid (1119-62-6) were obtained from Sigma Aldrich, U.S.A.. Ethyl acetate (141-78-6), n-hexane (110-54-3), Methanol (67-56-1) and Hydrogen peroxide (000-37655) were obtained from KISHIDA CHEMICAL, Japan. Azidoacetic acid NHS ester (824426-32-6), Cy5 acid (1032678-07-1) and DiSulfo-Cy3 azide (2055138-89-9) were obtained from Broadpharm, U.S.A.. 4- (tert-Butoxycarbonylamino)-1-butanol (75178-87-9) was obtained from Combi Blocks, U.S.A.. Boc-NH-PEG10k-NHS (Boc-PEG-NHS, PEG1110, Average molecular weight: 10 kDa) was obtained from Iris Biotech GmbH, Germany. Poly (glutamic acid) (PGA; 26247-79-0. Average molecular weight: 120 kDa) was obtained from Alamanda Polymers, U.S.A.. DMT-MM・$nH_2O$ (2170798-10-2) was obtained from WATANABE CHEMICAL, Japan. Human Carbonic Anhydrase IX (38-414), Fc, Avitag (CA9-H82F5) (CA9-H82F5-25 μg) was obtained from ACROBiosystems, China. Human breast cancer cell line (MDA-MB-231, EC92020424-F0) was obtained from KAC Co., Ltd, Japan.

**Instrumentation**

The reaction progress was monitored on TLC Silica gel 60 F254 (Sigma Aldrich, U.S.A.) and spots were visualized under 254 nm UV light. Column chromatography was performed using Wakosil 60, 64 ~ 210μm (Wako, Japan) as the stationary phase. Polymers were purified by dialysis using Visking Tube (For dialysis, cellulose material, MWCO: 3500 Da) (AS ONE,

Japan) and Spectra/Por® 7 Dialysis Membrane (MWCO: 15 kDa) (Repligen, U.S.A.). $^{1}H$ NMR spectra were corrected on JNM-GSX 400 (JEOL, Japan). UV-Vis, circular dichroism and fluorescence spectra were recorded using UV-Visible Spectrophotometer V-670, Spectrofluorometer FP-8500, Circular dichroism spectrometer J-725 (JASCO, Japan), respectively. UV-Vis spectra were also recorded NanoDrop 2000c (Thermo Fisher Scientific, U.S.A.) for small volume-samples. Dynamic light scattering measurement were performed using Disposable low-volume cuvette ZEN0040 and Zetasizer Nano ZS (Malvern, U.S.A). Solution pH was measured by LAQUA twin pH-22B (Horiba, Japan). Transmission electron microscopy observation was performed using carbon coated Cu grid (Nissin-EM, Japan) and TEM microscope H-7000 (Hitachi, Japan). SAXS measurement was performed with a lab-source small-angle instrument SAXSpace (Anton Paar, Austria) equipped with a 1D hybrid pixel detector Mythen 1K (Dectris, Switzerland) and Peltier-controlled sample holder TCStage 150 (Anton Paar, Austria). SAXS analysis were conducted by SAXSanalysis software (Anton Paar, Austria). Quartz crystal microbalance measurement was performed using AFFINIX Q8 and AFFINIX Q8 Sensor QCM01S (Piezo Parts Co.,Ltd., Japan). UV-vis and fluorescence measurements of low-volume samples were carried out using Synergy HTX (WakenBtech Co.,Ltd, Japan) and SH-9000Lab (Hitachi, Japan). Confocal images were taken using FV3000 (Olympus, Japan), respectively. Confocal images were analyzed using Imaris Cell Imaging Software 9.0 (Bitplane, UK) and Fiji [2].

### 2. Synthesis of CAIX inhibitor U-104

**Scheme S-1.** Synthesis of CAIX inhibitor U-104

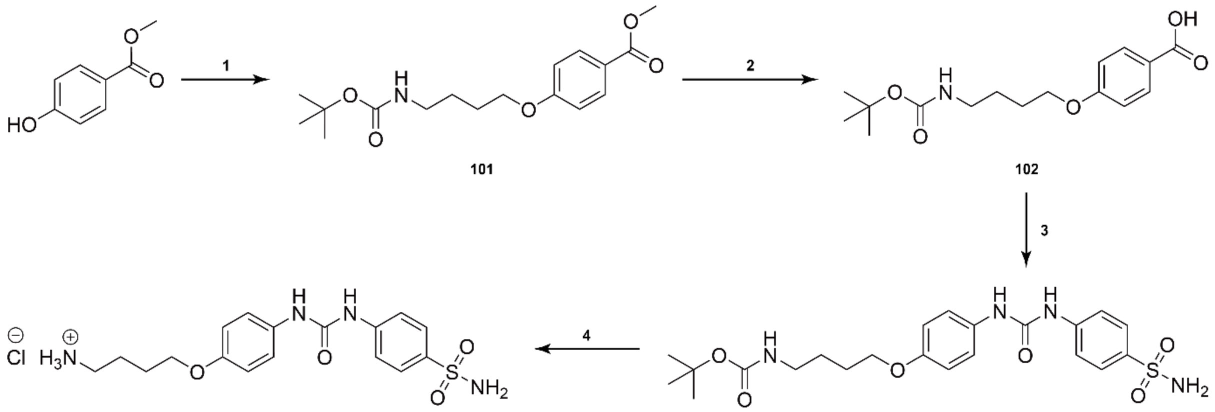


CAIX inhibitor U-104 was synthesized as previously described in the literature.[1]

#### 2-1. Synthesis of compound 101

Methyl 4-hydroxybenzoate (5.37 g, 35.3 mmol) and triphenylphosphine (13.9 g, 52.9 mmol) were dissolved in THF dehydrated (210 mL). Then, 4- (tert-Butoxycarbonylamino)-1-butanol (10.0 g, 52.9 mmol) dissolved in THF dehydrated (10 mL) was added into the solution. Diisopropyl azodicarboxylate (10.7 g, 52.9 mmol) was dropped to the reaction solution. The reaction was stirred overnight under a nitrogen atmosphere at room temperature. The reaction mixture was evaporated to remove solvent. Ethyl acetate was added, and supernatant was purified by column chromatography using hexane and ethyl acetate (v/v, 3:1, Rf = 0.37) and evaporated to yield a white powder (8.54 g, 50%). $^1$H NMR (400 MHz, $CDCl_3$): δ 7.97 (d, J = 12 Hz, 2H), 6.89 (d, J = 9.2 Hz, 2H), 4.60 (s, 1H), 4.03 (t, J = 6.4 Hz, 2H), 3.88 (s, 3H), 3.20 (m, 2H), 1.87-1.80 (m, 2H), 1.71-1.64 (m, 2H), 1.44 (s, 9H).

**2-2. Synthesis of compound 102**

**101** (4.55 g, 14.1 mmol) was dissolved in methanol (84.7 mL) and 5 M NaOH aq. (84.7 mL) was added. The reaction mixture was left stirring for 3 h at 50 °C. The reaction was traced by thin-layer chromatography (TLC) analysis using hexane and ethyl acetate (*v*/*v*, 3:1, Rf of compound **102** = 0). The reaction mixture was evaporated to remove methanol, then, diluted with ethyl acetate (250 mL). The aqueous phase was acidified with 10% $KHSO_4$ and extracted twice with ethyl acetate (100 mL). We checked the existence of compound **102** in the water phase with TLC using hexane and ethyl acetate (*v*/*v*, 3:1) but there was no spot showing compound **102**. Then, the organic phase was evaporated and dried in a vacuum to yield a white powder (3.21 g, 88%). $^1$H NMR (400 MHz, $CDCl_3$): δ 8.03 (d, J = 9.2 Hz, 2H), 6.92 (d, J = 8.8 Hz, 2H), 4.61 (s, 1H), 4.05 (t, J = 5.6 Hz, 2H), 3.21 (m, 2H), 1.88-1.81 (m, 2H), 1.72-1.65 (m, 2H), 1.45 (s, 9H).

**2-3. Synthesis of compound 103**

**102** (3.21 g, 10.39 mmol) was dissolved in toluene dehydrates (65 mL), and *N, N*-diisopropylethylamine (2.71 g, 3.57 mL, 21.00 mmol) was added under a nitrogen atmosphere at 90 °C. Under the same condition, diphenyl phosphoryl azide (3.18 g, 11.56 mmol) was added to the reaction mixture in one portion and the mixture was stirred for 5 h. The reaction was traced by TLC analysis using hexane and ethyl acetate (*v*/*v*, 2:5, Rf of the product = 0.35). The reaction mixture was then evaporated and dissolved in acetonitrile hydrated (50 mL), then, the mixture was heated to 60 °C. Sulfanilamide (2.71 g, 15.76 mmol, Rf = 0.36) was added in one portion, then, the reaction mixture was stirred overnight under a nitrogen atmosphere. The reaction mixture was evaporated and dried in a vacuum for 5 h. The crude product was dissolved in ethyl acetate (50 mL), washed 20 times with 30 mM HCl aq. Then the organic layer was evaporated to obtain brown precipitation. The precipitation was purified by column chromatography using hexane and ethyl acetate (v/v, 2:5, Rf of compound **103**= 0.25) and evaporated to yield a white powder. The crude product was washed by benzene and dried in a vacuum to yield a white powder (838.7 mg, 17%). $^{1}$H NMR (400 MHz, DMSO-*d6*): δ 8.99 (s, 1H), 8.60 (s, 1H), 7.71 (d, J = 8.8 Hz, 2H), 7.59 (d, J = 9.2 Hz, 2H), 7.34 (d, J = 8.8 Hz, 2H), 7.19 (s, 2H), 6.86 (d, J = 9.2 Hz, 3H), 3.91 (t, J = 6.4 Hz, 2H), 2.96 (q, J = 6.3 Hz, 2H), 1.70-1.63 (m, 2H), 1.51 (q, J = 7.7 Hz, 2H), 1.37 (s, 9H).

#### 2-4. Synthesis of U-104

Under a nitrogen atmosphere, add 5 mL of THF (dehydrated) to a weighed amount of 103 (0.765 mg, 1.60 mmol), sonicate to dissolve, and add 6.5 mL of HCl/ethyl acetate. Stir for about 120 minutes to allow the reaction to proceed. The HCl/ethyl acetate is removed by nitrogen flow. No purification is performed. The reaction is followed by TLC using hexane and ethyl acetate (v/v, 2:5, Rf of compound **103**= 0.25). The powder was dried in vacuum to obtain a white powder (581 mg, 17%). $^1$H NMR (400 MHz, $D_2O$): δ 7.84 (d, J = 9.1 Hz, 2H), 7.55 (d, J = 9.1 Hz, 2H), 7.29 (d, J = 9.1 Hz, 2H), 7.00 (d, J = 9.1 Hz, 2H), 4.09 (t, J = 5.4 Hz, 2H), 7.19 (s, 2H), 6.86 (d, J = 9.2 Hz, 3H), 3.91 (t, J = 6.4 Hz, 2H), 3.05 (t, J = 6.3 Hz, 2H), 1.86-1.81 (m, 4H).

### 3. Synthesis of $N_3$-PEG-U

**Scheme S-2.** Synthesis route of **$N_3$-PEG-U**

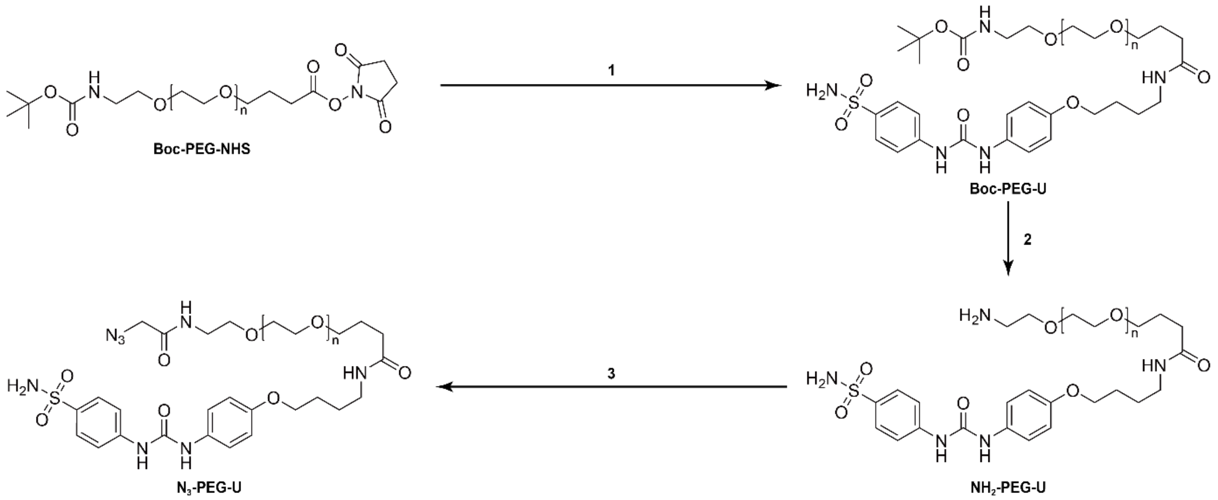


#### 3-1. Synthesis of Boc-PEG-U

To a solution of U-104 (41.6 mg, 0.100 mmol) and Triethylamine (67.6 µg, 0.400 mmol) in dry DMF (2 mL) was added Boc-PEG-NHS (100.0 mg, 0.0100 mmol). The mixture was stirred at 37 °C for 24 h. The reaction mixtures were dialyzed on a dialysis membrane (MWCO: 3500, regenerated cellulose) against milli-Q for 2 days, followed by lyophilized to obtain Boc-PEG-U (98.6 mg, 96%). The grafting degree of U-104 was calculated from $^1$H NMR spectrum.

#### 3-2. Synthesis of $NH_2$-PEG-U

To a solution of Boc-PEG-U (169.7 mg, 0.0162 mmol) in milli-Q (4.0 mL) water was added 5 M HCl aq. (80.0 μL). The mixture was stirred at 37 °C for 48 h. The reaction mixtures were dialyzed on a dialysis membrane (MWCO: 3500, regenerated cellulose) against milli-Q for 1 day, followed by lyophilized to obtain $NH_2$-PEG-U (148.6 mg, 89%). Deprotection rate of *N*-Boc group was calculated from $^1$H NMR spectrum.

#### 3-3. Synthesis of $N_3$-PEG-U

To a solution of Triethylamine (35.3 μL, 0.200 mmol) in dry DMF (0.5 mL) was added $NH_2$-PEG-U (50.0 mg, 0.00489 mmol). The mixture was stirred and a solution of Azidoacetic acid NHS ester (9.9 mg, 0.0500 mmol) in dry DMF (0.5 mL) was added dropwise. The mixture was stirred at 37 °C for 48 h. The reaction mixtures were dialyzed on a dialysis membrane (MWCO: 3500, regenerated cellulose) against milli-Q for 2 days, followed by lyophilized to obtain $N_3$-PEG-U (43.5 mg, 86%). Grafting degree of azide group was calculated from $^1$H NMR spectrum.

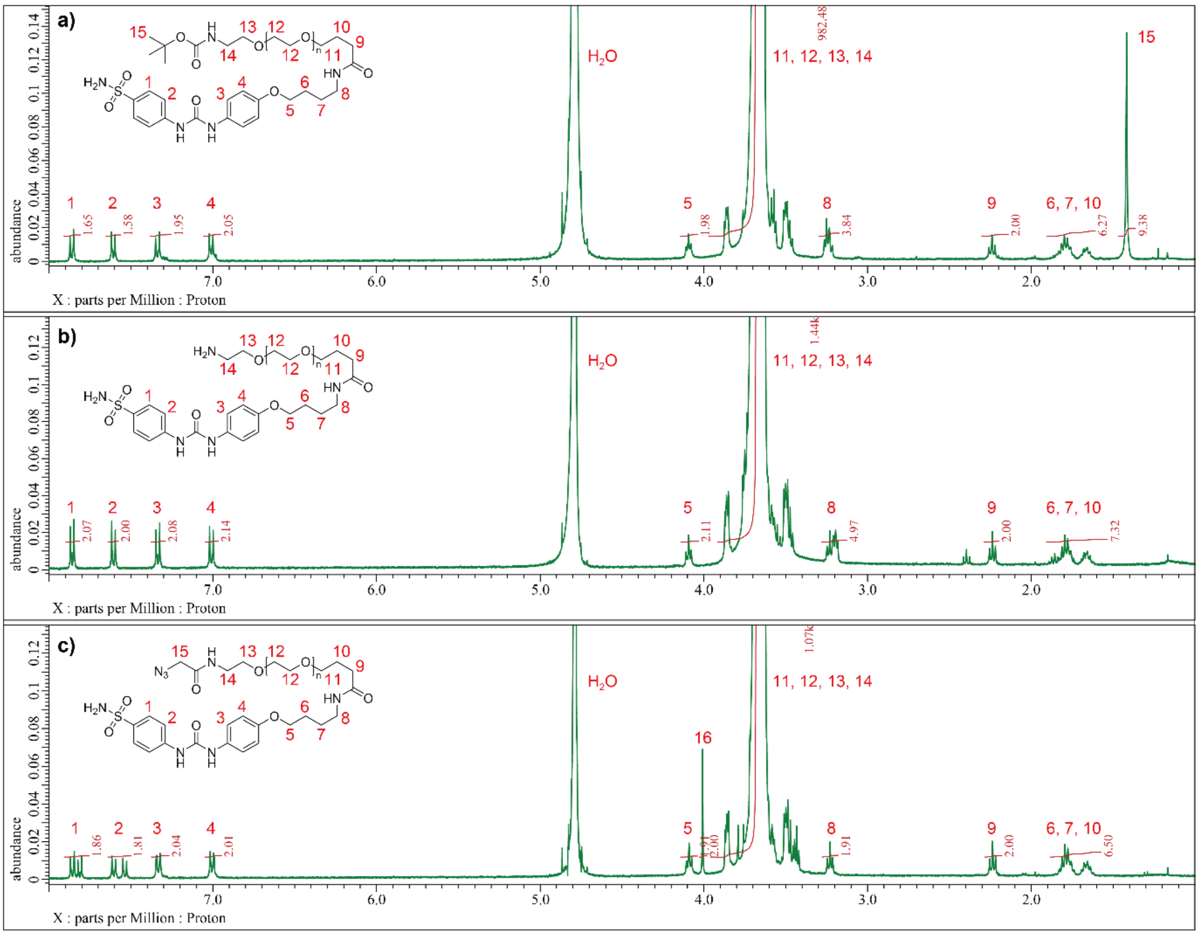


**Figure S-1.** $^1$H NMR spectra of (a) Boc-PEG-U, (b) $NH_2$-PEG-U and (c) $N_3$-PEG-U (400 MHz, 25 °C). The grafting degree of U moiety and Azide group were calculated from the integration of the peak 15 and 16, respectively.

### 4. Evaluation of physico-chemical properties for $N_3$-PEG-U

#### 4-1. Filter experiment

$N_3$-PEG-U (0.5 mg, 0.05 mmol) and compound 103 (0.12 mg, 0.251 mmol) were dissolved in D-PBS (1 and 5 mL), respectively, and shaken at room temperature for 5 min. Samples were centrifuged (10000 G, 5 min, r.t.), and UV-vis measurement was performed for the supernatant using NanoDrop 2000c.

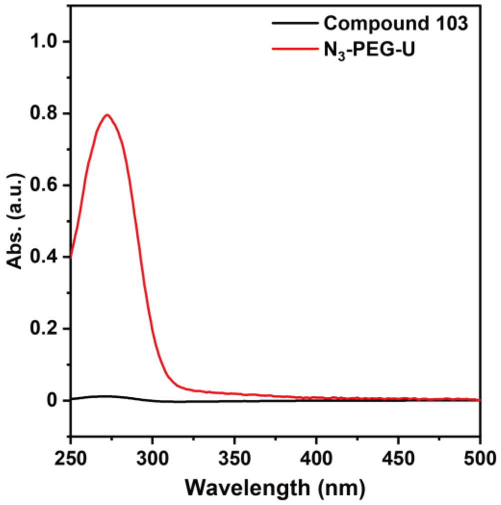


**Figure S-2.** UV-vis spectra of compound 103 (black) and $N_3$-PEG-U (red).

#### 4-2. Size evaluation by dynamic light scattering (DLS)

For DLS measurements, a solution of $N_3$-PEG-U series in D-PBS (0.1 mL) was dropped into the solution and injected into a disposable low-volume cuvette. The DLS measurements of the polymers were all performed on a Zetasizer NanoZS with a 633 nm laser (4 mW). Samples were equilibrated for 2 min before analysis and more than 3 runs per sample were taken. Besides, the scattering angle was fixed at 173°.

#### 4-3. Circular dichroism (CD) measurement for $N_3$-PEG-U

$N_3$-PEG-U (1.0 mg, 0.10 mmol) was dissolved in D-PBS and methanol (1 mL), respectively. CD spectra of $N_3$-PEG-U solution was monitored using circular dichroism spectrometer J-725. A cuvette with 0.1 mm path length was used, and the photomultiplier voltage did not exceed 600 V in this measurement.

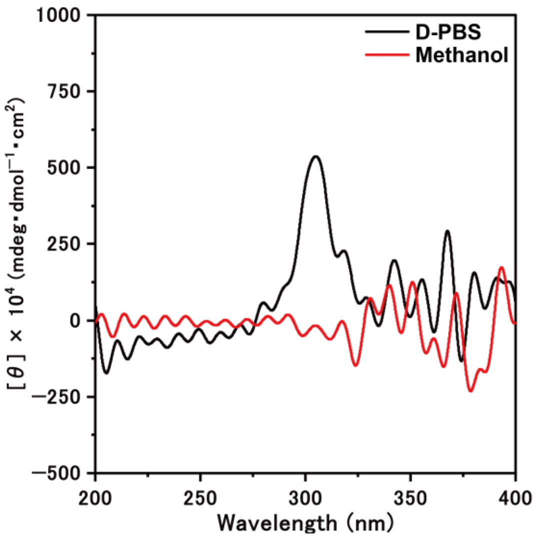


**Figure S-3.** CD spectra of $N_3$-PEG-U in D-PBS (black) and methanol (red).

#### 4-4. Determination of Critical self-assemble concentration (CSC) for $N_3$-PEG-U by pyrene fluorescence titrations

A series of polymer solutions in milli-Q at various concentrations were prepared. A stock solution of pyrene (5.2 mg $L^{-1}$ in Acetonitrile, 23 μL) was added to a solution of $N_3$-PEG-U (1 mL). The resulting solution was sonicated for 30 min at 4 °C. Subsequently, the solvent was removed under reduced pressure. D-PBS (1 mL) was added to the prepared thin film and the solution was incubated overnight at room temperature. The fluorescence of each pyrene-containing $N_3$-PEG-U solution (having different concentrations) was measured at an excitation wavelength of 334 nm using spectrofluorometer FP-8500 through a 1 × 1 cm quartz cell. CSC was determined by analyzing the fluorescence intensities of solubilized pyrene's first and third vibronic band maxima $I_I$ and $I_{III}$ at 374 nm and 384 nm. These ratios $I_I$ / $I_{III}$ were plotted against log polymer concentration and linear extrapolation; the CSC was determined by calculating the break point of $I_I/I_{III}$ based on the piecewise regression shown in equation (1) and (2).

$$I_{373}/I_{384} = \frac{I_{373}/I_{384_1}(\mathrm{LogCSC}-\mathrm{LogC})+I_{373}/I_{384_{CSC}}(\mathrm{LogC}-\mathrm{LogC_1})}{\mathrm{LogCSC}-\mathrm{LogC_1}} \; if \; \mathrm{LogC} < \mathrm{LogCSC} \quad (1)$$

$$I_{373}/I_{384} = \frac{I_{373}/I_{384_{CSC}}(\mathrm{LogC_2}-\mathrm{LogC})+I_{373}/I_{384_2}(\mathrm{LogC}-\mathrm{LogCSC})}{\mathrm{LogC_1}-\mathrm{LogCSC}} \; if \; \mathrm{LogC} > \mathrm{LogCSC} \quad (2)$$

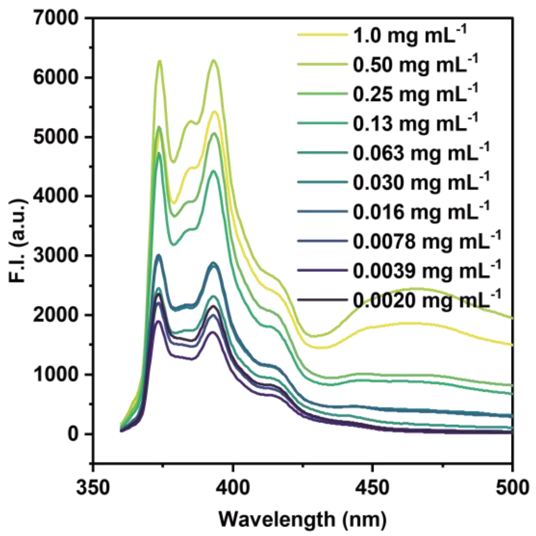


**Figure S-4.** Fluorescence spectra of $N_3$-PEG-U solution at various concentrations.

### 4-5. Determination of CSC for $N_3$-PEG-U by DLS

For DLS measurements, a solution of $N_3$-PEG-U in D-PBS (0.1 mL) at various concentrations was dropped into the solution and injected into a disposable low-volume cuvette. The DLS measurements of the polymers were all performed on a Zetasizer NanoZS with a 633 nm laser (4 mW). Samples were equilibrated for 2 min before analysis and more than 3 runs per sample were taken. Besides, the scattering angle was fixed at 173°. These logarithmic scattering intensities were plotted against logarithmic $N_3$-PEG-U concentration; the CSC was determined by calculating the break point based on the piecewise regression shown in equation (3) and (4).

$$\mathrm{LogInt} = \frac{\mathrm{Int_1(LogCSC - LogC) + Int_{CAC}(LogC - LogC_1)}}{\mathrm{LogCSC - LogC_1}} \text{ if } \mathrm{LogC} < \mathrm{LogCSC} \quad (3)$$

$$\mathrm{LogInt} = \frac{\mathrm{Int_{CSC}(LogC_2 - LogC) + Int_2(LogC - LogCSC)}}{\mathrm{LogC_1 - LogCSC}} \text{ if } \mathrm{LogC} > \mathrm{LogCSC} \quad (4)$$

### 5. Synthesis of DBCO-PGA

#### 5-1. Synthesis of DBCO-PGA

100 mM carbonate-bicarbonate buffer was prepared by mixing sodium carbonate (53.0 mg, 0.500 mmol) and sodium hydrogen carbonate (43.4 mg, 0.516 mmol) in milli-Q (10 mL), 0.1 M HCl aq. was added to adjust the pH to the 8.45-8.54 range. Poly (glutamic acid) (49.6 mg, 0.413 μmol) was dissolved in 22.4 mM carbonate-bicarbonate buffer (14.9 mL) and sonicated for 10 min. A solution of DMT-MM (50.1 mg, 0.167 mmol) in milli-Q (0.1 mL) was dropped into the solution and stirred for 10 min. A solution of DBCO-amine (9.12 mg, 0.0330 mmol) in THF (3 mL) was added dropwise. The reaction mixture was stirred at room temperature for 24 h. The reaction mixtures were dialyzed on a dialysis membrane (MWCO: 120000, regenerated cellulose) against milli-Q and MeOH for 3 days (Day 1 and 3: milli-Q, Day 2: MeOH), followed by lyophilized to obtain DBCO-PGA (24.1 m, 49%). Grafting degree of DBCO group was calculated from $^{1}H$ NMR spectrum.

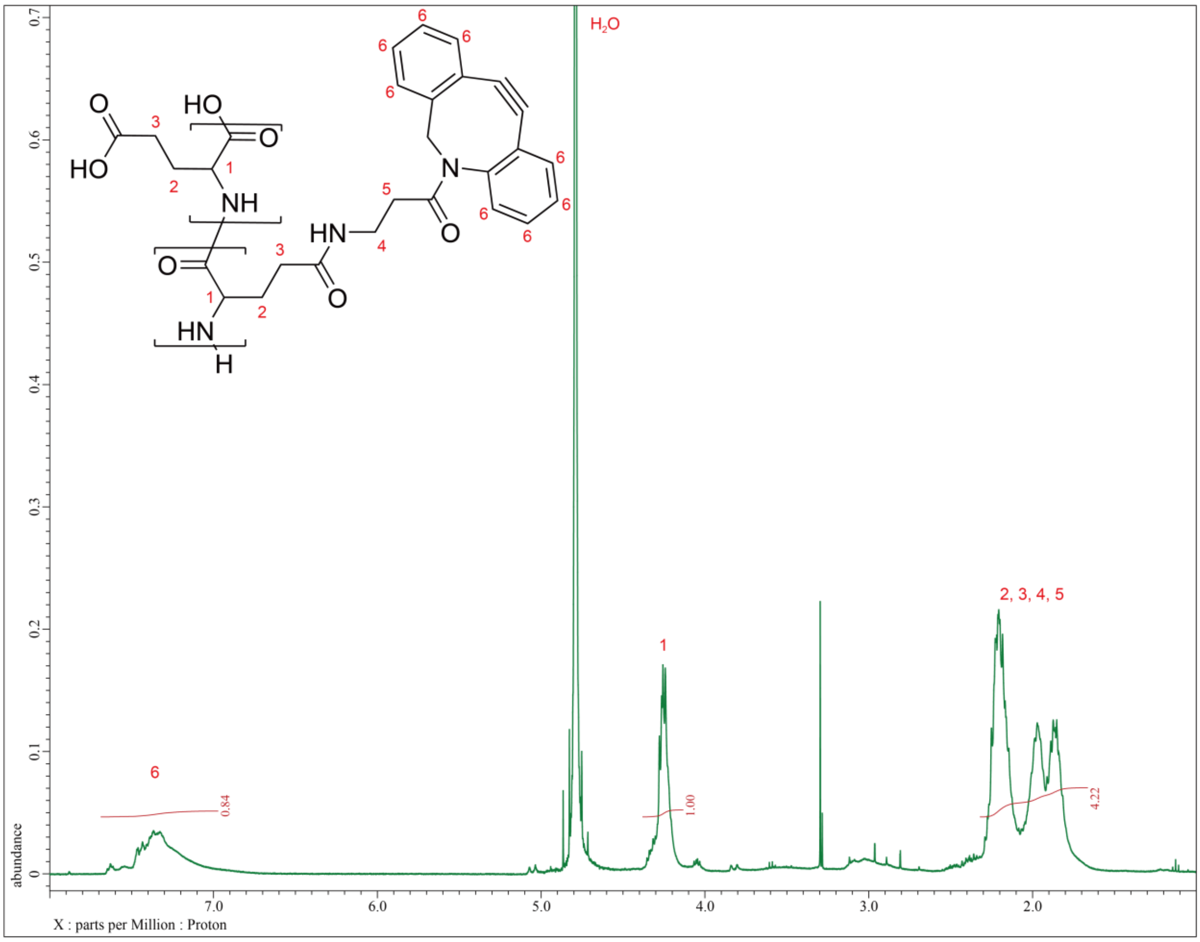


**Figure S-5.** $^{1}H$ NMR spectra of DBCO-PGA (400 MHz, 10 mM NaOD, 25 °C). The grafting degree of DBCO group was calculated from the integration of the peak 1 and 6.

### 6. Evaluation of physico-chemical properties for DBCO-PGA

#### 6-1. Filter experiment

DBCO-PGA (0.375 mg, 3.13 nmol) and DBCO-acetate (0.080 mg, 0.251 μmol) was dissolved in D-PBS (5 mL), respectively. Shaken at room temperature for 5 min. Samples were centrifuged (10000 G, 5 min, r.t.) then UV-vis measurement was performed for the supernatant using NanoDrop 2000c.

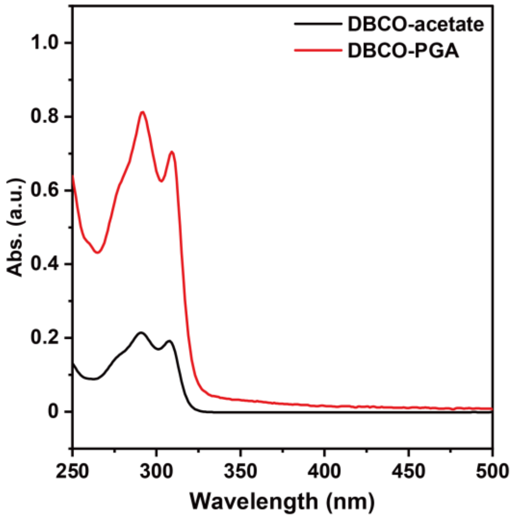


**Figure S-6.** UV-vis spectra of DBCO-acetate (black) and DBCO-PGA (red).

#### 6-2. Size evaluation by dynamic light scattering (DLS)

For DLS measurements, a solution of DBCO-PGA series in D-PBS (0.1 mL) was dropped into the solution injected into a disposable low volume cuvette. The DLS measurement of the polymers were all performed on a Zetasizer NanoZS instrument with a 633 nm laser (4 mW). Samples were equilibrated for 2 min before analysis and more than 3 runs per sample taken. Besides, the scattering angle was fixed at 173°.

#### 6-3. CSC for DBCO-PGA by pyrene fluorescence titrations

A series of polymer solutions in milli-Q at various concentrations were prepared. A stock solution of pyrene (5.2 mg $L^{-1}$ in Acetonitrile, 23 μL) was added to a solution of DBCO-PGA (1 mL). The resulting solution was sonicated for 30 min at 4 °C. Subsequently, the solvent was removed under reduced pressure. D-PBS (1 mL) was added to the prepared thin film and the solution was incubated overnight at room temperature. The fluorescence of each pyrene-containing DBCO-PGA solution (having different concentrations) was measured at an excitation wavelength of 334 nm using a fluorimeter (RF-6000, Shimadzu) through a 1 × 1 cm quartz cell. CSC was determined by analyzing the fluorescence intensities of solubilized pyrene's first and third vibronic band maxima $I_I$ and $I_{III}$ at 374 nm and 384 nm. These ratios $I_I$ / $I_{III}$ were plotted against log polymer concentration and linear extrapolation; the CSC was determined by calculating the break point of $I_I/I_{III}$ based on the piecewise regression shown in equation (1) and (2).

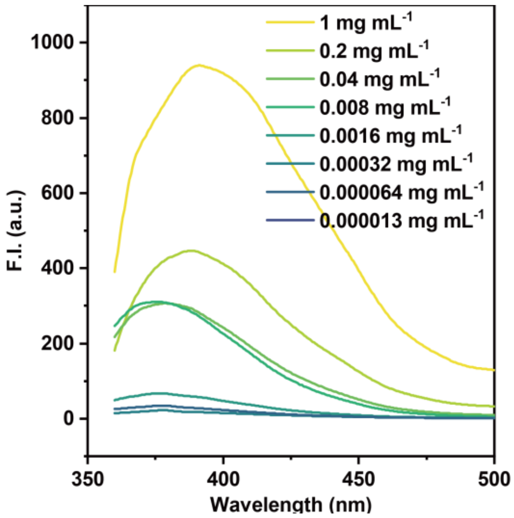


**Figure S-7.** Fluorescence spectra of DBCO-PGA solution at various concentrations.

#### 6-4. Determination of CSC for DBCO-PGA by DLS

For DLS measurements, a solution of DBCO-PGA in D-PBS (0.1 mL) at various concentrations was dropped into the solution and injected into a disposable low-volume cuvette. The DLS measurements of the polymers were all performed on a Zetasizer NanoZS instrument with a 633 nm laser (4 mW). Samples were equilibrated for 2 min before analysis and more than 3 runs per sample were taken. Besides, the scattering angle was fixed at 173°. These logarithmic scattering intensities were plotted against logarithmic DBCO-PGA concentration; the CSC was determined by calculating the break point based on the piecewise regression shown in equation (3) and (4).

### 7. Change of physico-chemical properties of nano-assemblies via click reaction

#### 7-1. Evaluation of the stability of nano-assemblies by DLS

For DLS measurements, a solution of polymers in D-PBS (0.1 mL) was dropped into the solution injected into a disposable low volume cuvette (ZEN0040, Malvern, Germany). The DLS measurement of the polymers were all performed on a Malvern Zetasizer NanoZS instrument with a 633 nm laser (4 mW). Samples were equilibrated for 2 min before analysis and 3 runs per sample taken every 30 min. Besides, the scattering angle was fixed at 173°.

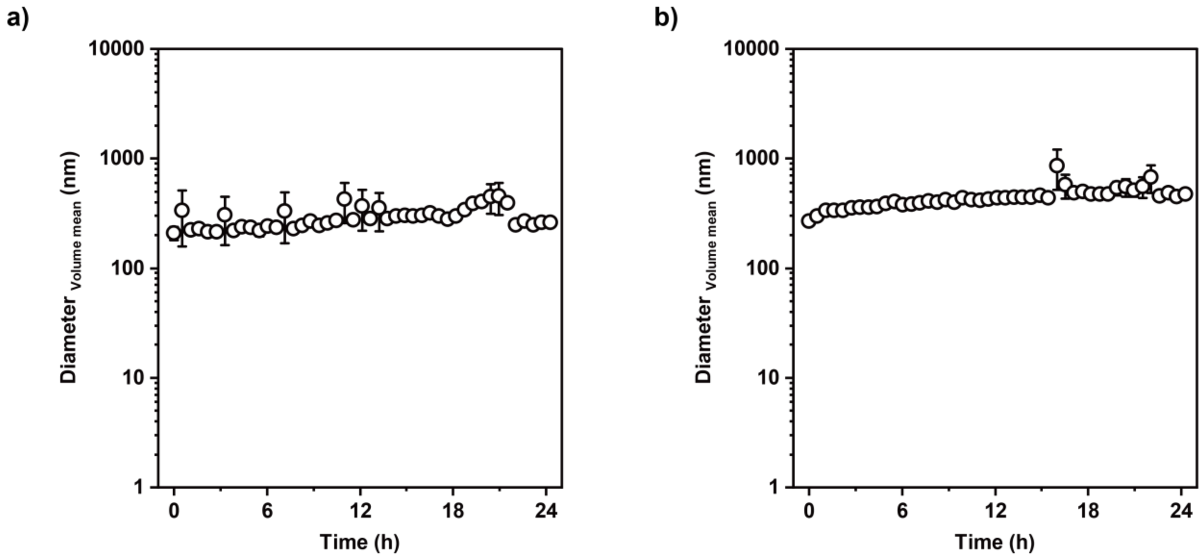


**Figure S-8.** Time-course of the volume-averaged diameter of a) $N_3$-PEG-U and b) DBCO-PGA (1.0 mg $mL^{-1}$).

### 7-2. Checking the conversion of click reaction under DLS conditions.

DBCO-PGA (0.6 mg, 0.4 mmol) and $N_3$-PEG-U (3.0 mg, 0.30 mmol) were dissolved in D-PBS (1 mL). DBCO-PGA soln. (50 μL) was added to $N_3$-PEG-U soln. (50 μL), then the mixture soln. UV-vis measurements were performed immediately after mixing and after 6 hours.

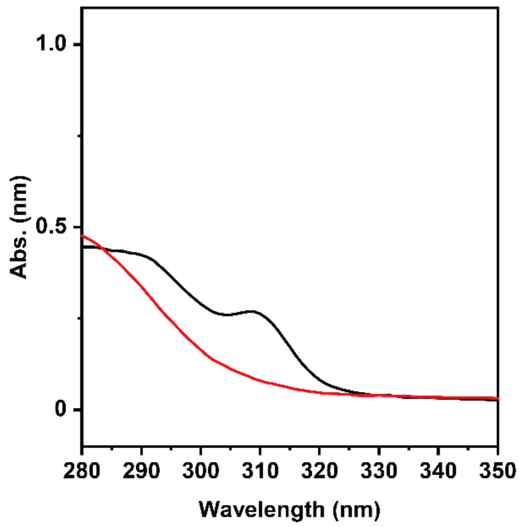


**Figure S-9.** UV-vis spectra of samples immediately after mixing and after 6 hours.

**7-3. Size evaluation after click reaction by DLS**

DBCO-PGA (0.3 mg, 0.2 mmol) and/or $N_3$-PEG-U (1.5 mg, 0.15 mmol) were dissolved in D-PBS (1 mL). The solution was injected into a disposable low volume cuvette. Samples were equilibrated for 2 min before analysis and 1 run per sample taken. Besides, the scattering angle was fixed at 173°.

**7-4. Evaluation of size changes during click reaction**

For DLS measurements, a solution of polymers in D-PBS (0.1 mL) was dropped into the solution injected into a disposable low volume cuvette (ZEN0040, Malvern, Germany). The DLS measurement of the polymers were all performed on a Malvern Zetasizer NanoZS instrument with a 633 nm laser (4 mW). Samples were equilibrated for 2 min before analysis and more than 3 runs per sample taken. Besides, the scattering angle was fixed at 173°. DLS measurement was repeated over 300 times every 5 sec.

**7-5. Determination of Critical self-assemble concentration (CSC) for PGA-trz-PEG-U by pyrene fluorescence titrations**

A series of polymer solutions in milli-Q at various concentrations were prepared. A stock solution of pyrene (5.2 mg $L^{-1}$ in Acetonitrile, 23 μL) was added to a solution of PGA-trz-PEG-U (1 mL). The resulting solution was sonicated for 30 min at 4 °C. Subsequently, the solvent was removed under reduced pressure. D-PBS (1 mL) was added to the prepared thin film and the solution was incubated overnight at room temperature. The fluorescence of each pyrene-containing PGA-trz-PEG-U solution (having different concentrations) was measured at an excitation wavelength of 334 nm using spectrofluorometer FP-8500 through a 1 × 1 cm quartz cell. CSC was determined by analyzing the fluorescence intensities of solubilized pyrene's first and third vibronic band maxima $I_{I}$ and $I_{III}$ at 374 nm and 384 nm. These ratios $I_{I}$ / $I_{III}$ were plotted against log polymer concentration and linear extrapolation.

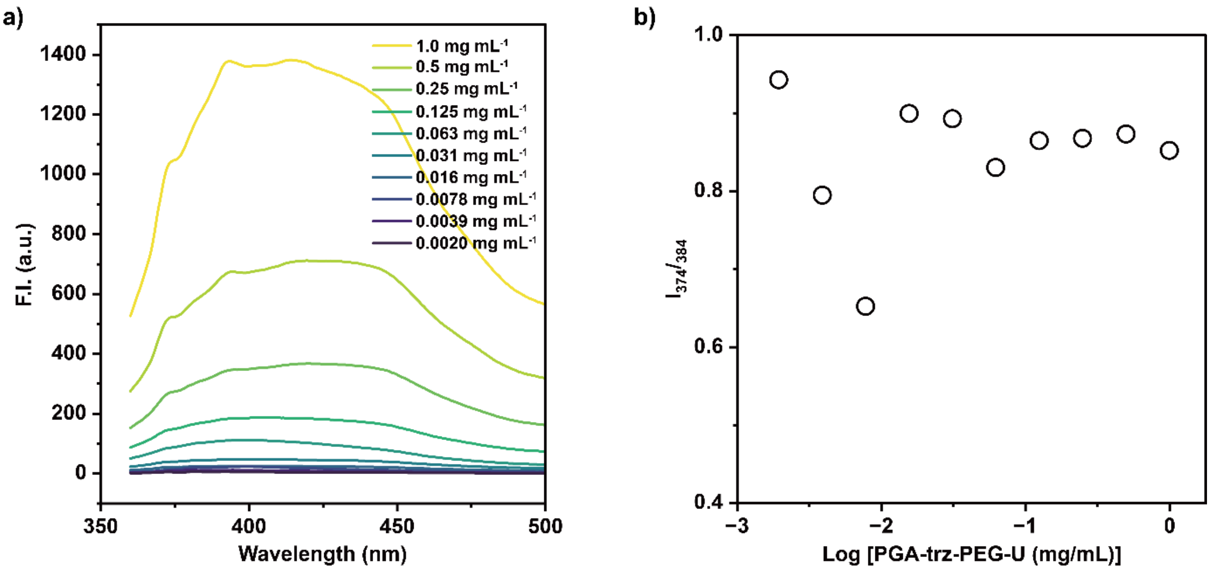


**Figure S-10.** a) Fluorescence spectra of PGA-trz-PEG-U solution at various concentrations. b) CSC determination of multivalent ligands: PGA-trz-PEG-U.

#### 7-6. Small Angle X-ray Scattering (SAXS) experiments

SAXS was conducted with a lab-source small-angle instrument SAXSpace equipped with a 1D hybrid pixel detector Mythen 1K using line-collimated Cu Kα radiation (1.54 Å). The samples were loaded in a 1-mm quartz capillary cell and the sample temperature was controlled by using a Peltier-controlled sample holder TCStage 150. Scattering curves of $N_3$-PEG-U, DBCO-PGA, and buffer were acquired at 25 ◦C, and the in-situ scattering curves during the click-reaction at 1 h, 3 h, and 6 h were acquired at 37 °C in vacuum conditions, respectively. The sample-to-detector distance was 317 mm. The exposure time per sample was 1 h. For all experiments, the attenuated primary beam at q = 0 was monitored by using a semitransparent beam stop. SAXS patterns were normalized with the primary-beam intensity to unity by using SAXSanalysis software. Background subtraction (capillary and corresponding buffer) and collimation correction (desmearing) were also performed by SAXSanalysis software [3].

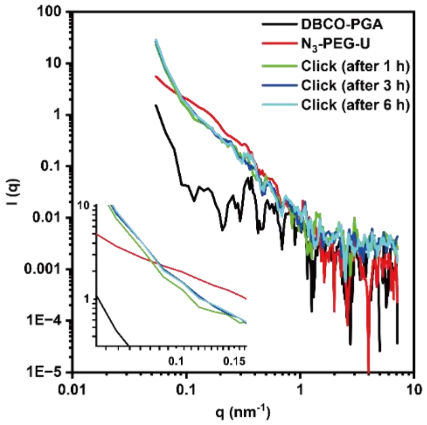


**Figure S-11.** SAXS profile of DBCO-PGA and/or $N_3$-PEG-U in D-PBS as solvent. The inset shows the magnified view of the low-q region, emphasizing the changes in particle size. Note that time-dependent increase in intensity at 0.1 $nm^{-1}$ supports the conformational change.

#### 7-7. Transmission electron microscopy (TEM) observation

One drop of a solution of polymers was dropcasted on carbon coated Cu grid (Nissin-EM, Tokyo, Japan) and lyophilized overnight. The samples obtained were stained with 2% uranyl acetate before observation using a TEM microscope (H-7000, Hitachi, Tokyo, Japan).

#### 7-8. Calculation of Circularity

The circularity defined by equation (5) was used to evaluate the morphology of the structures, where A is the area of the structure and p is the perimeter. Circularity was determined by manually analyzing the TEM images using ImageJ (more than 30).

$$\text{Circularity} = \frac{4\pi A}{\text{p}} \quad (5)$$

### 8. Reaction kinetics evaluation of click reaction

#### 8-1. Monitoring click reaction

DBCO-PGA (0.3 mg, 0.2 mmol) and DBCO-amine (1 mg, 3.62 mmol) was dissolved in D-PBS (2 mL) and THF (1 mL), respectively. Then, the concentration of alkyne solution was adjusted to 100 mM based on standard curve by UV-vis spectroscopy. $N_3$-PEG-U (2 mg, 0.2 mmol) and azide acetic acid (10 mL, 0.133 mmol) was dissolved in D-PBS (2 and 1 mL), respectively. In a quartz cuvette equipped, 1.8 mL of 100 mM alkyne solution was added, followed by 1.8 mL of 100 mM azide solution. The cuvette was thermostated at 37 ºC with a water-circulating thermostat. UV-vis measurement (190-800 nm) was repeated over 24 hours every 30 min.

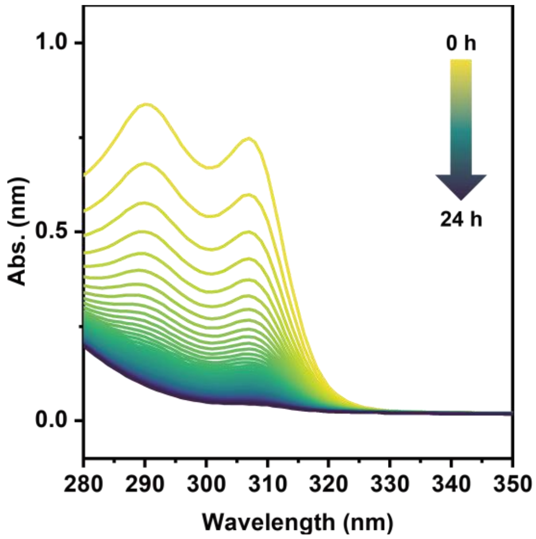


**Figure S-12.** UV-vis spectra with click reaction between $N_3$-AcOH and DBCO-amine.

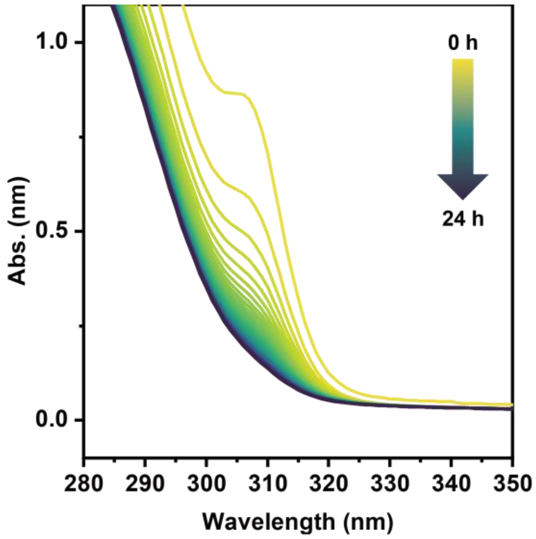


**Figure S-13.** UV-vis spectra with click reaction between $N_3$-PEG-U and DBCO-amine.

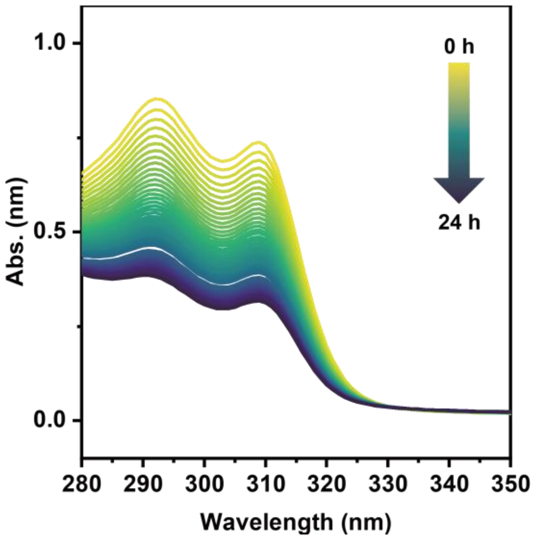


**Figure S-14.** UV-vis spectra with click reaction between $N_3$-AcOH and DBCO-PGA.

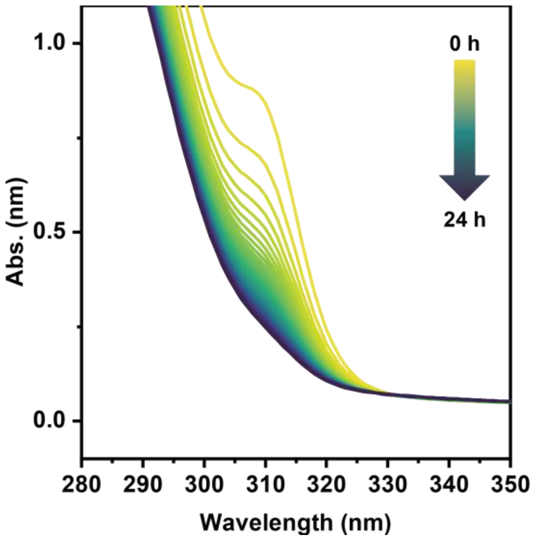


**Figure S-15.** UV-vis spectra with click reaction between $N_3$-PEG-U and DBCO-PGA.

### 8-2. Kinetic analysis

The absorbance at 309 nm in each of the spectra (Figure S-10 - S-13) obtained in 8-1. is plotted against time. For samples which contain $N_3$-PEG-U, the absorbance at 309 nm of the same concentration of $N_3$-PEG-U was subtracted from the plot to accommodate scattering. Rate constants were obtained by curve fitting using equation (6), which was derived from the rate equations for second-order reactions in the same molecule.

$$[A] = \frac{[A]_0}{1+kt[A]_0} \quad (6)$$

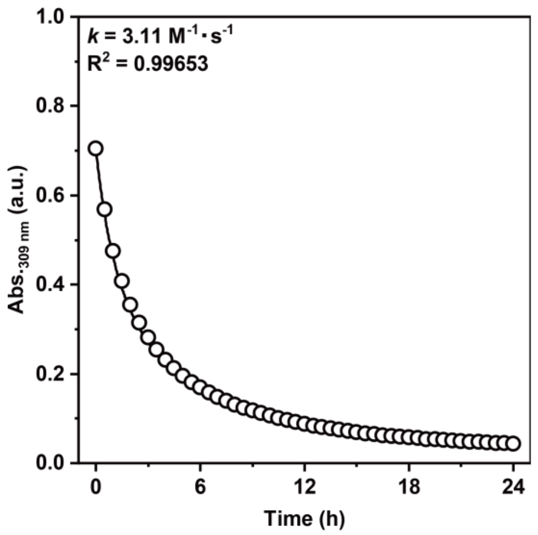


**Figure S-16.** Time course of absorbance at 309 nm with click reaction between $N_3$-AcOH and DBCO-amine.

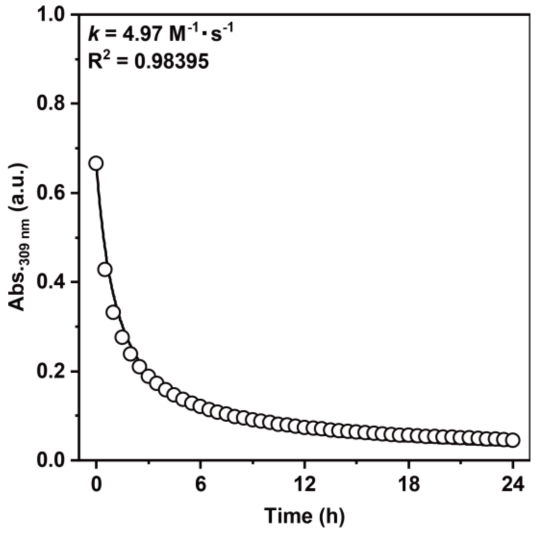


**Figure S-17.** Time course of absorbance at 309 nm with click reaction between $N_3$-PEG-U and DBCO-amine.

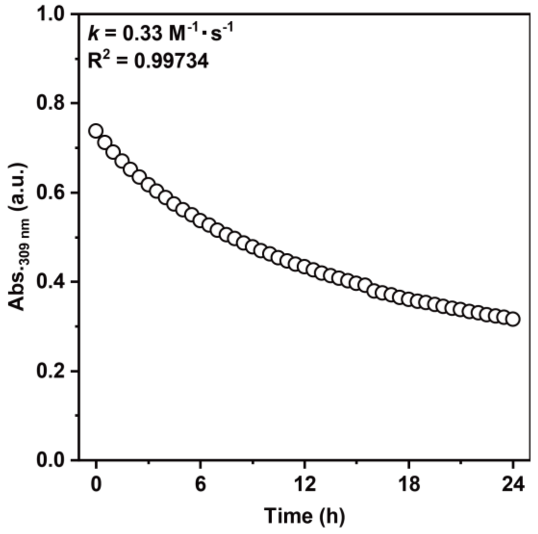


**Figure S-18.** Time course of absorbance at 309 nm with click reaction between $N_3$-AcOH and DBCO-PGA.

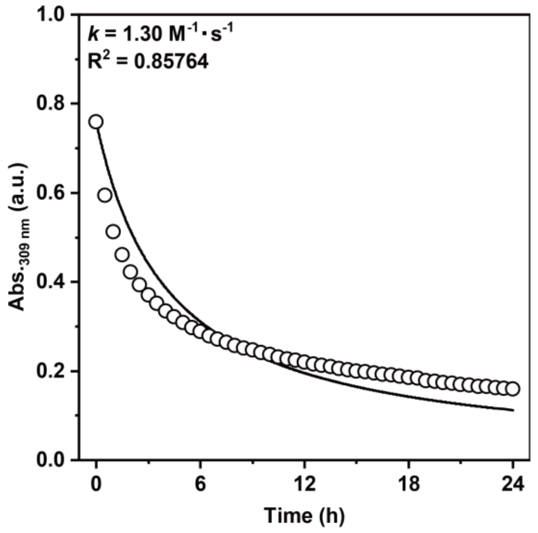


**Figure S-19.** Time course of absorbance at 309 nm with click reaction between $N_3$-PEG-U and DBCO-PGA.

### 8-3. Curve-fitting for the calculation of the conversion

Table S-1. The fitting parameter and $R^2$ values of the fitting curve (equation (6)) for the calculation of the conversion.

| Substrate combination | $[A]_0$ (M) | $k$ ($M^{-1}$ $h^{-1}$) | Adj. $R^2$ |
|---|---|---|---|
| $N_3$-AcOH and DBCO-amine | 9.97E-05 | 7259.64499 | 0.9591 |
| $N_3$-PEG-U and DBCO-amine | 9.41E-05 | 14209.61354 | 0.98677 |
| $N_3$-AcOH and DBCO-PGA | 9.99E-05 | 723.43862 | 0.99965 |
| $N_3$-PEG-U and DBCO-PGA | 9.82E-05 | 4335.69653 | 0.981 |

Fitting equation: $\text{Conversion (\%)} = 100 \times \frac{C_0kt}{1+C_0kt}$ (6)

### 9. Binding assay by quartz crystal microbalance (QCM)

#### 9-1. QCM measurement

A 27 MHz QCM system was used to monitor and quantify interactions between the polymers and carbonic anhydrase IX (CAIX). All QCM experiment in this study were conducted at 25 °C.

#### 9-2. Immobilization of Streptavidin (SAv) on the surface QCM sensors

SAv was immobilized on the QCM electrode via biotin-avidin recognition. First, gold electrodes were cleaned three times with piranha solution (fresh mixture of $H_2O_2$ (aq.) and $H_2SO_4$; 30% $H_2O_2$ (aq.)/ $H_2SO_4$ = 1:3 (*v*/*v*)) for 5 min. Next, 3,3′-dithiodipropionic acid (5 µL, 20 mM in EtOH) was loaded into the QCM cells and then incubated for more than 30 min. The resulting cells were washed with milli-Q and then the carboxylic acid groups on the gold surface were activated for 30 min by loading 100 µL of a 1:1 *v*/*v* aqueous solution of 1-ethyl-3-(3-dimethylaminopropyl) carbodiimide (100 $mg\cdot mL^{-1}$) and N-hydroxy succinimide (100 $mg\cdot mL^{-1}$) to form N-hydroxy succinimide (NHS) esters. After rinsing the activated cells with milli-Q three times, D-PBS was added to the cells. The amine solution (mixture of 33 mM Biotin-PEG2-amine (aq.) and 33 mM 2-amino ethanol (aq.); Biotin-PEG2-amine: 2-amino ethanol = 20:80 (*v*/*v*)) was loaded into the activated cells and then incubated for more than 30 min. After rinsing the biotinylated cells with milli-Q several times, D-PBS was added to the cells and the cells were set on the QCM system. After stabilization of the oscillation frequency,

SAv (concentration in the cell after injection: 1 μM) was injected on the biotinylated QCM electrodes and then incubated for more than 30 min.

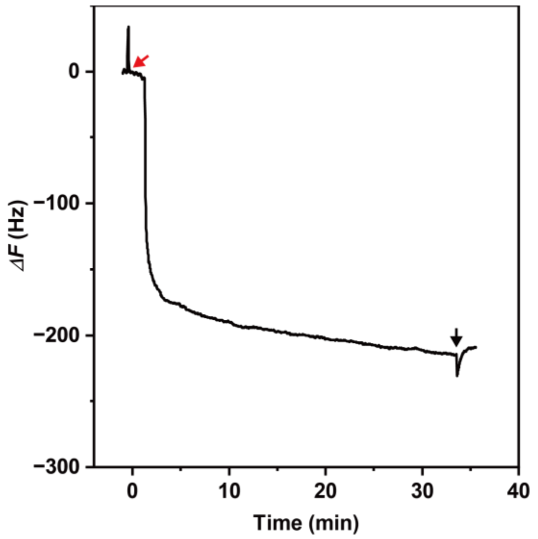


**Figure S-20.** Time courses of QCM frequency change with SAv injection (red arrow) and wash by D-PBS (black arrow).

**9-3. Immobilization of CAIX on QCM sensors**

CAIX was immobilized on the QCM electrode via a biotin-avidin recognition. After rinsing the SAv-immobilized cells to remove the non-immobilized SAv using D-PBS, the cells were filled with the buffer (100 μL). After stabilization of the oscillation frequency, Biotinylated Human Carbonic Anhydrase IX (38-414), Fc, Avitag (CA9-H82F5) (CAIX, concentration in the cell after injection: 50 nM) was injected on the SAv-immobilized cells and then incubated for more than 30 min.

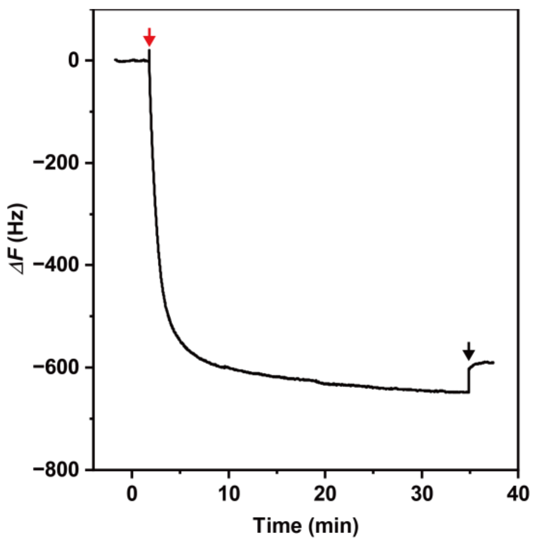


**Figure S-21.** Time courses of QCM frequency change with CAIX injection (red arrow) and wash by D-PBS (black arrow).

#### 9-4. Immobilization of Casein on QCM sensors

After rinsing the CAIX-immobilized cells to remove the non-immobilized CAIX using D-PBS, the cells were filled with the buffer (100 μL). After stabilization of the oscillation frequency, Casein (concentration in the cell after injection: 0.01 w/v%) was injected on the CAIX-immobilized cells and then incubated for more than 30 min.

#### 9-5. Synthesis of PGA-trz-PEG-U with the multivalency of 70

DBCO-PGA (0.3 mg, 0.2 mmol) was dissolved in D-PBS (2 mL). $N_3$-PEG-U (1 mg, 0.1 mmol) was dissolved in D-PBS (1 mL), respectively. $N_3$-PEG-U soln. and DBCO-PGA soln. were added in 1 mL each to the eppendorf tubes, pipetted and then incubated at 37 ºC for 20 h.

The reaction mixture was diluted using D-PBS and then used as PGA-trz-PEG-U with multivalency of 70 for binding assays using QCM.

#### 9-6. Immobilization of polymers on CAIX-immobilized QCM sensors

Frequency change was measured for CAIX-immobilized QCM sensors when solutions of $N_3$-PEG-U and DBCO-PGA and PGA-trz-PEG-U were added sequentially every 30 min, and curve fitting was performed based on the Langmuir model (equation (7)). PGA-trz-PEG-U was synthesized by incubating $N_3$-PEG-U (50 μM) and DBCO-PGA ([DBCO] = (50 μM)) at 37 °C for 20 h.

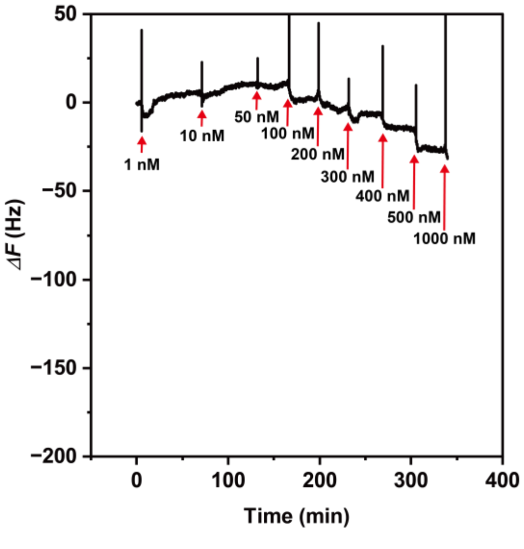


**Figure S-22.** Time courses of QCM frequency change with $N_3$-PEG-U injection (red arrow).

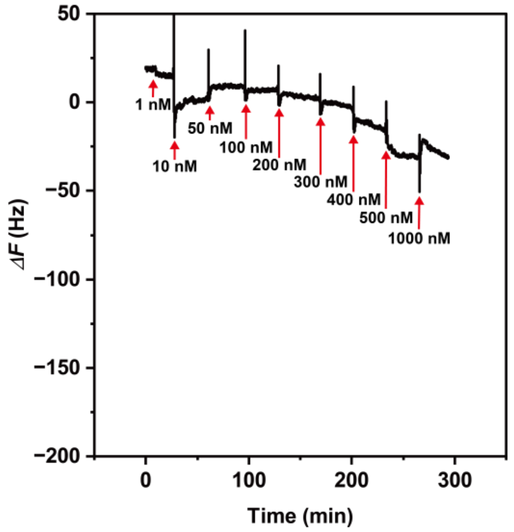


**Figure S-22.** Time courses of QCM frequency change with DBCO-PGA injection (red arrow).

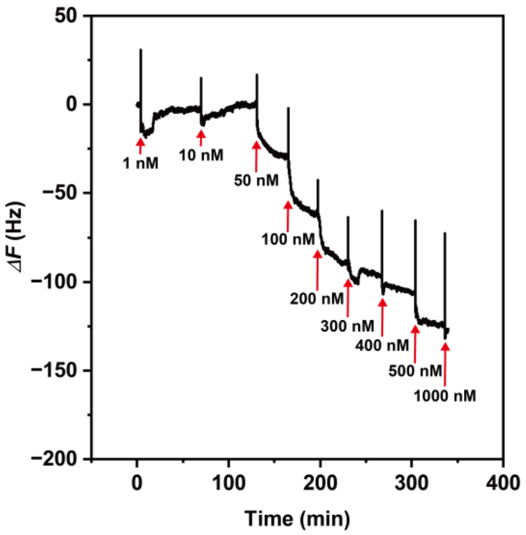


**Figure S-24.** Time courses of QCM frequency change with PGA-trz-PEG-U injection (red arrow).

$$-\Delta F = \frac{-\Delta F_{max} \times [PGA-trz-PEG-U]_0}{K_d + [PGA-trz-PEG-U]_0} \quad (7)$$

**9-7. Synthesis of PGA-trz-PEG-U with the multivalency of 15 and 30**

DBCO-PGA (0.3 mg, 0.2 mmol) was dissolved in D-PBS (2 mL). $N_3$-PEG-U (1 mg, 0.1 mmol) was dissolved in D-PBS (1 mL), respectively. $N_3$-PEG-U soln. and DBCO-PGA soln. and D-PBS were added to Eppendorf tubes in volume ratios (0: 1: 1) or (0.5: 1: 0.5) or (0.25: 1: 0.75), pipetted and incubated at 37°C for 20 hours. The reaction mixture was diluted using D-PBS and then used as PGA-trz-PEG-U with multivalency of 0 or 15 or 30 for binding assays using QCM. In addition, the progress of click reaction was monitored by UV-vis spectroscopy to calculate the conversion rate after 20 hours.

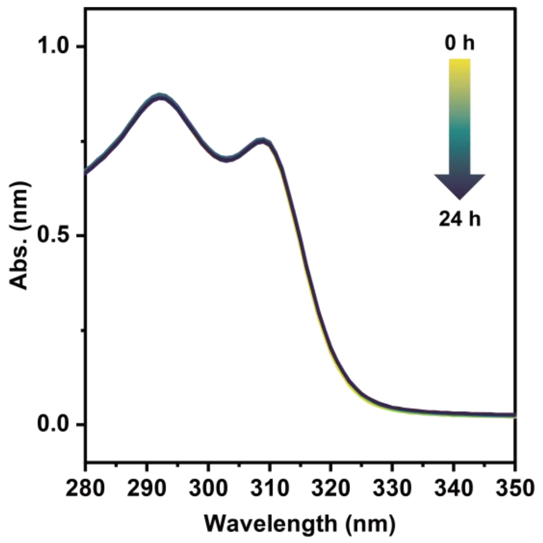


**Figure S-25.** UV-vis spectra of DBCO-PGA in D-PBS for 24 h ([DBCO] = 50 μM).

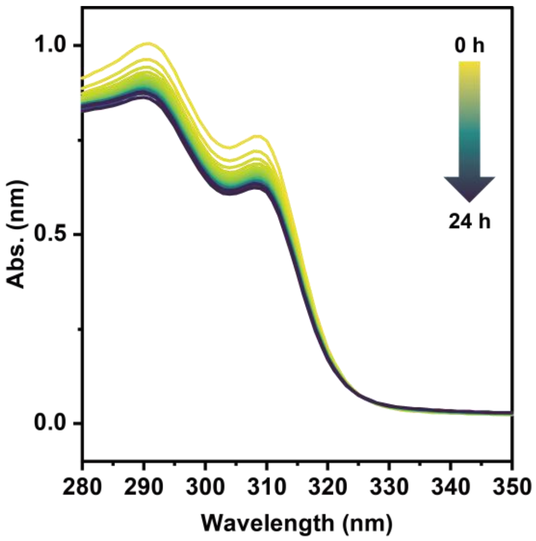


**Figure S-26.** UV-vis spectra with click reaction between $N_3$-PEG-U and DBCO-PGA ([$N_3$] = 12.5 μM and [DBCO] = 50 μM).

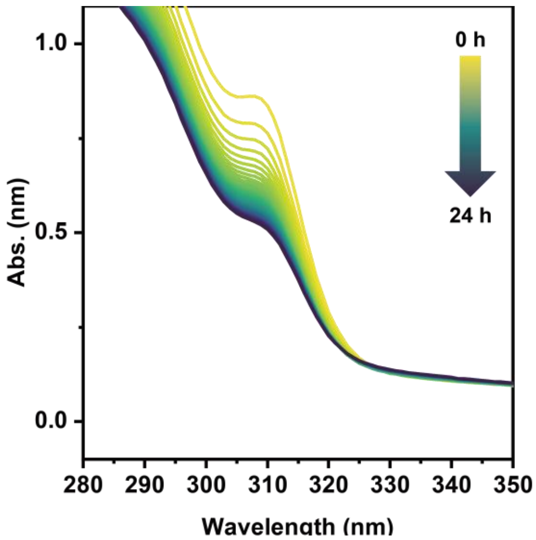


**Figure S-27.** UV-vis spectra with click reaction between $N_3$-PEG-U and DBCO-PGA ([$N_3$] = 25 μM and [DBCO] = 50 μM).

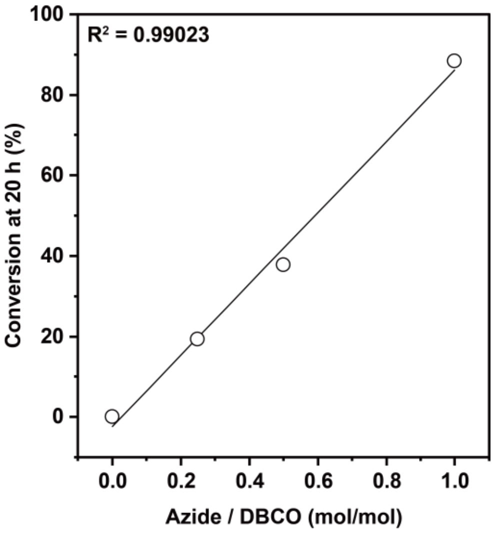


**Figure S-28.** Relationship between $N_3$ and DBCO ratios and conversion rates after 20 hours.

### 10. Cell experiments

#### 10-1. Cell culture

Human breast cancer cell line (MDA-MB-231) was obtained from KAC Co., Ltd. (Kyoto, Japan, EC92020424-F0). MDA-MB-231 cells were cultured in DMEM supplemented with 10% FBS and 1% antibiotics. For culture under hypoxic conditions, cells were incubated at 37 °C under an atmosphere of 1% $O_2$, 5% $CO_2$, and 94% $N_2$ in a humidified incubator. For culture under normoxic conditions, cells were grown in a humidified incubator at 37 °C under 5% $CO_2$.

#### 10-2. Water-Soluble Tetrazolium Salts Assay

The water-soluble tetrazolium salt assay (WST assay) was performed to determine cell viability treated with polymers. MDA-MB-231 cells ($5.0 \times 10^3$ cell/well) were seeded onto 96-well plates using DMEM adjusted to pH 6.8 or 7.4 using 1 M HCl aq. and incubated for 24 h under 1% $O_2$ (hypoxia) or normoxia. After incubation, cells were treated with polymers and incubated for 72 h under hypoxic or normoxic conditions. Then, the plates were washed with D-PBS, and cell counting kit-8 solution diluted 10 times by DMEM was added. After 3 h, the plates were diluted 2 times by D-PBS and centrifuged (4000 rpm, 5 min). Then, 80 µL of the supernatant was collected to 96-well plates, and the absorbance at 450 nm was measured using a microplate reader.

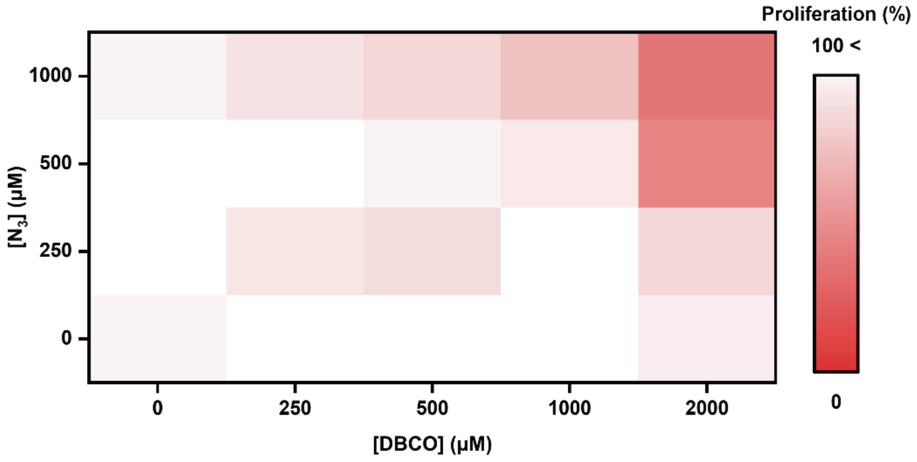


**Figure S-29.** Heatmap depicting the proliferative inhibition of MDA-MB-231 by the combination of polymers.

### 10-3. Checking the progress of click reaction in the media

DBCO-PGA (3.0 mg, 0.025 μmol) was dissolved in DMEM (500 μL). $N_3$-PEG-U (1 mg, 0.10 μmol) was also dissolved in DMEM (50 μL). $N_3$-PEG-U soln. and DBCO-PGA soln. were added in 20 μL each to the eppendorf tubes, pipetted and then incubated at 37 °C. The reaction mixture was diluted 10-fold using milli-Q and then UV-vis measurements were performed.

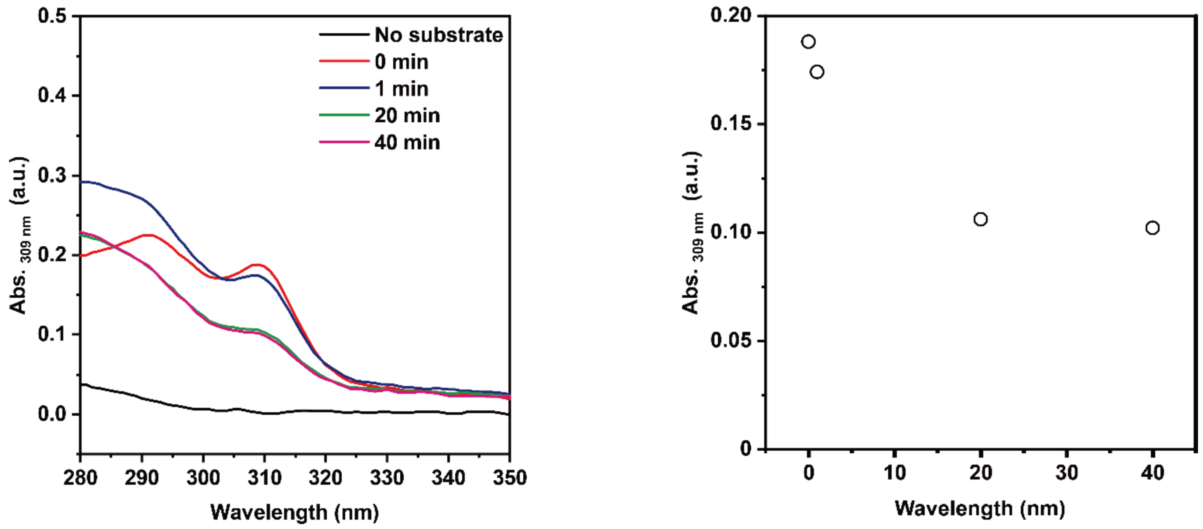


**Figure S-30.** UV-visible spectra following click reaction in DMEM.

### 10-4. Pre-synthesized/purified of PGA-trz-PEG-U for the evaluation of the cell guided effect

DBCO-PGA (1.2 mg, 0.010 μmol) was dissolved in D-PBS (296 μL). $N_3$-PEG-U (4.0 mg, 0.40 μmol) was also dissolved in D-PBS (296 μL). The solution of DBCO-PGA was added the solution of $N_3$-PEG-U and incubated at 37 °C. After 11 h, the sample was diluted 10-fold and then UV-vis measurements were performed. After checking the progress of click reaction by UV-vis spectoscopy, PGA-trz-PEG-U was obtained by lyophilization.

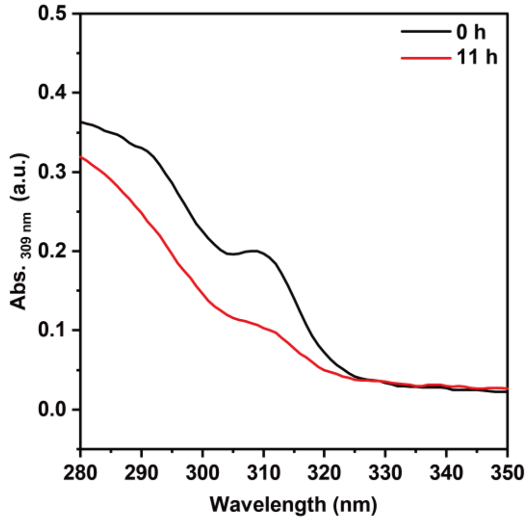


**Figure S-31.** UV-visible spectra before and after click reaction in the pre-synthesis of PGA-trz-PEG-U.

#### 10-4. Synthesis of fluorescence-labeled $N_3$-PEG-U ($N_3$-PEG-U-Cy5)

A solution of $N_3$-PEG-U (10 mg, 1.0 μmol) and Cy5 acid (2.56 mg, 5.0 μmol) in anhydrous DMF (200 μL) were added DMAP (6.11 mg, 50 μmol) and EDCI·HCl (9.6 mg, 50 μmol). After stirring at 37 °C for 24 hours., The reaction mixtures were dialyzed on a dialysis membrane (MWCO: 12-14 kDa, regenerated cellulose) against methanol and milli-Q for 2 days (Day 1: methanol, Day 2: milli-Q), followed by lyophilized to obtain $N_3$-PEG-U-Cy5 (8.0 mg, 80%). Grafting degree of Cy5 group was calculated using calibration line by fluorescence measurement.

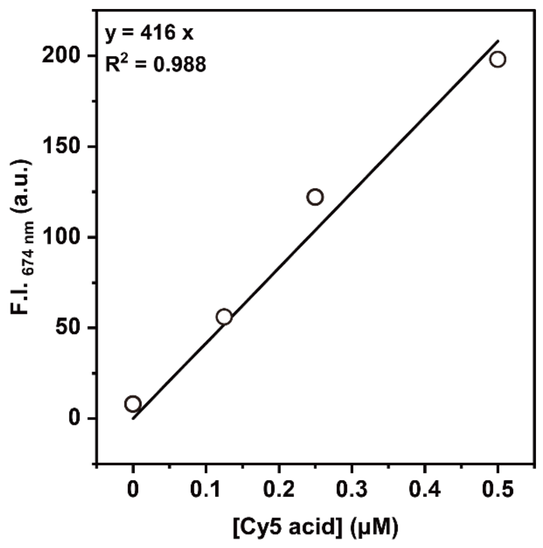


**Figure S-32.** Calibration line of Cy5 acid in D-PBS.

### 10-5. Synthesis of fluorescence-labeled DBCO-PGA (DBCO-PGA-Cy3)

DBCO-PGA (5 mg, 3.34 μmol) was dissolved in D-PBS (5 mL). Disulfo-Cy3-azide (250 μg, 0.335 μmol) was added to the solution and stirred at 37 °C for 24 hours. The reaction mixtures were dialyzed on a dialysis membrane (MWCO: 12-14 kDa, regenerated cellulose) against milli-Q water for 2 days, followed by lyophilized to obtain DBCO-PGA-Cy3 (4.4 mg, 88%). Grafting degree of Cy3 group was calculated using calibration line by fluorescence measurement.

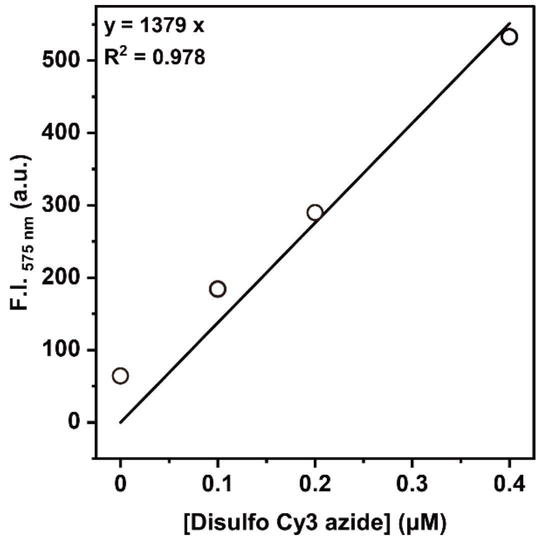


**Figure S-33.** Calibration line of disulfo Cy3 azide in D-PBS.

#### 10-6. Visualization of the Binding of polymers on MDA-MB-231

The binding of polymers on MDA-MB-231 was visualized using fluorescence-labeled $N_3$-PEG-U ($N_3$-PEG-U-Cy5) and fluorescence-labeled DBCO-PGA (DBCO-PGA-Cy3) by confocal microscopy. MDA-MB-231 ($1.5 \times 10^4$ cell/well) were grown in a 96-well glass bottom plate using DMEM adjusted to pH 6.8 by 1 M HCl aq. and incubated for 72 h in 1% $O_2$ (hypoxia). After incubation, $N_3$-PEG-U-Cy5 (final concentration of azide group =40 or 200 μM) and DBCO-PGA-Cy3 (final concentration of DBCO group = 80 or 400 μM) were added, and cells were incubated for 1 h at 37 °C. The medium was removed, and after washing (3 × 100 μL DMEM), Hoechst 33342 (1 μg $mL^{-1}$) was used to stain cell nuclei for 10 min. Confocal images were obtained with an FV3000 confocal microscope and taken using a 405 nm laser diode (0.3% or 0.6%) for excitation with emission collected from 430 to 470 nm (detector voltage 900 V or 740 V) for Hoechst 33342, a 640 nm laser diode (1.0% or 0.4%) for excitation with emission collected from 650 to 750 nm (detector voltage 680 V or 520 V) for Cy5, a 561 nm laser diode (0.3%) for excitation with emission collected from 570 nm to 670 nm (detector voltage 470 V or 420 V) for Cy3 and a 561 nm laser diode (0.5% or 0.1%) for excitation with emission collected from 654 nm or 640 nm to 752 nm and 694 nm (detector voltage 500 V or 520 V) for calculation of FRET ratio. Confocal images were processed and analyzed by Fiji to quantify the fluorescent intensities of polymers in cells. To quantify the fluorescent intensities, the images were preprocessed by the adjustment of brightness / contrast and applied median filter (1 pixel) in Fiji.

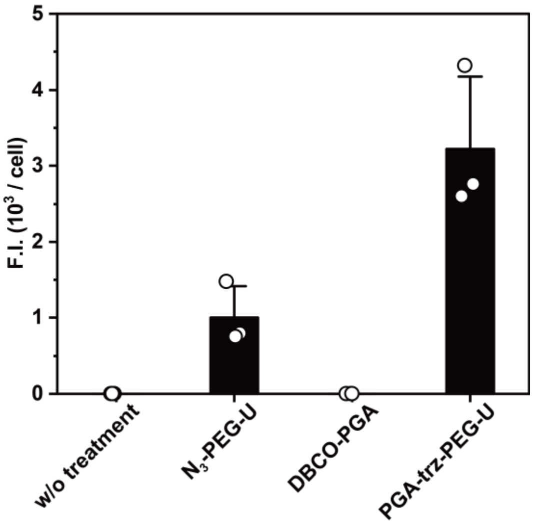


**Figure S-34.** The Cy5 fluorescent intensities excited by the excitation wavelength for Cy5 of cancer cells treated with $N_3$-PEG-U ([$N_3$] = 40 μM) and/or DBCO-PGA ([DBCO] = 80 μM).

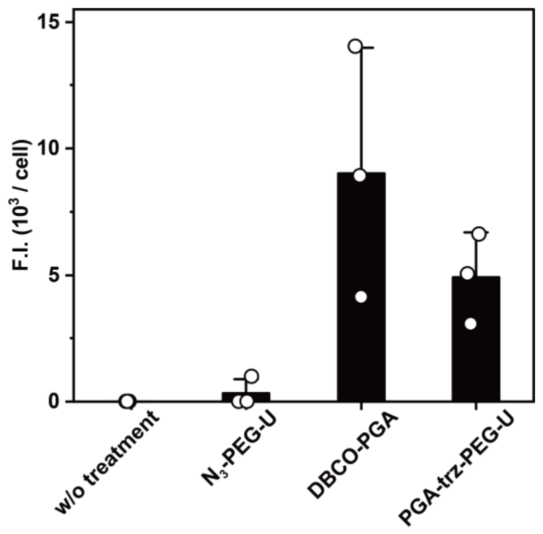


**Figure S-35.** The Cy3 fluorescent intensities excited by the excitation wavelength for Cy3 of cancer cells treated with $N_3$-PEG-U ($[N_3]$ = 40 μM) and/or DBCO-PGA ([DBCO] = 80 μM).

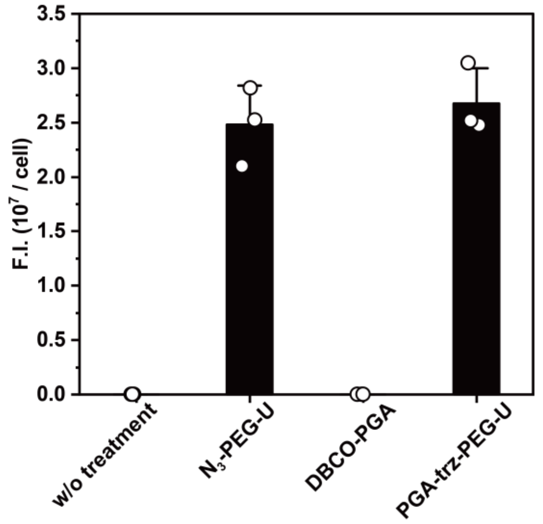


**Figure S-36.** The Cy5 fluorescent intensities excited by the excitation wavelength for Cy5 of cancer cells treated with $N_3$-PEG-U ($[N_3]$ = 200 μM) and/or DBCO-PGA ([DBCO] = 400 μM).

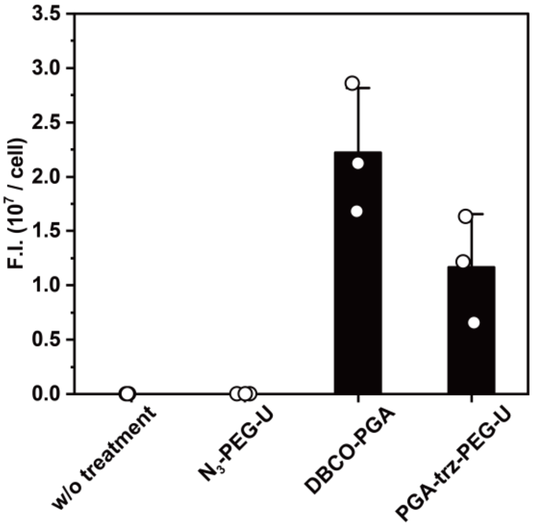


**Figure S-37.** The Cy5 fluorescent intensities excited by the excitation wavelength for Cy3 of cancer cells treated with $N_3$-PEG-U ([$N_3$] = 200 μM) and/or DBCO-PGA ([DBCO] = 400 μM).

### 10-7. Calculation of the FRET ratio

The FRET ratio was calculated by dividing the fluorescent intensities of Cy5 excited at the excitation wavelength for Cy3 ($F.I._{Cy5/Cy3}$) by the fluorescence intensity of Cy3 excited at the excitation wavelength for Cy3 ($F.I._{Cy3/Cy3}$) (equation (7)).

$$\text{FRET ratio} = \frac{\text{F.I.}_{Cy5/Cy3}}{\text{F.I.}_{Cy3/Cy3}} \quad (7)$$

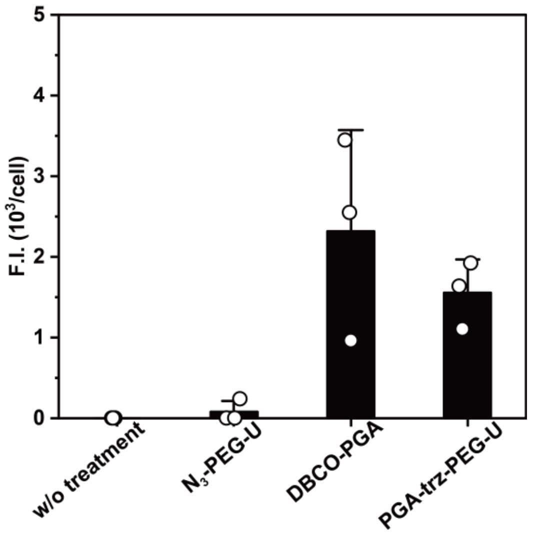


**Figure S-38.** The Cy5 fluorescent intensities excited by the excitation wavelength for Cy3 of cancer cells treated with $N_3$-PEG-U ([$N_3$] = 40 μM) and/or DBCO-PGA ([DBCO] = 80 μM).

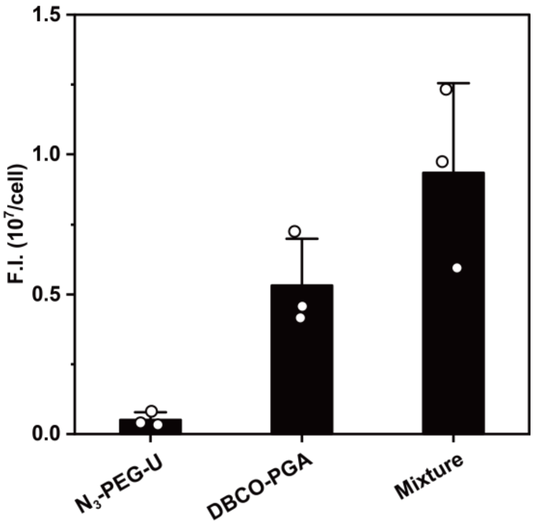


**Figure S-39.** The Cy5 fluorescent intensities excited by the excitation wavelength for Cy3 of cancer cells treated with $N_3$-PEG-U ([$N_3$] = 200 μM) and/or DBCO-PGA ([DBCO] = 400 μM).
